# Multi-epitope Peptide Vaccine Prediction for Candida albicans targeting Pyruvate Kinase Protein; an Immunoinformatics Approach

**DOI:** 10.1101/758920

**Authors:** Anfal Osama Mohamed Sati, Abdelrahman Hamza Abdelmoneim Hamza, Enas Dawoud Khairi Dawoud, Tebyan Ameer Abdelhameed Abbas, Fatima Abdelrahman Bshier Abdelrahman, Khalid Abbas Hassan Saad, Rouaa Babikir Ahmed Abduallah, Marwa Mohamed Elhag Saeed Mustafa, Tamador abdelrahman Sidahmed, Reham M. Elhassan, Mohamed A. Hassan

## Abstract

The fungus *Candida albicans* is an opportunistic pathogen that causes a wide range of infections. It’s the primary cause of candidiasis and the fourth most common cause of nosocomial infection. In addition, disseminated invasive candidiasis which is a major complication of the disease has an estimated mortality rate of 40%-60% even with the use of antifungal drugs. Over the last decades, several different anti-Candida vaccines have been suggested with different strategies for immunization against candidiasis such as, live-attenuated fungi, recombinant proteins, and glycoconjugates but none has been approved by the FDA, yet. This study aims to introduce a new possible vaccine for *C. albicans* through analyzing peptides of its pyruvate kinase (PK) protein as an immunogenic stimulant computationally.

A total number of 28 *C. albicans*, pyruvate kinase proteins were obtained from NCBI on the 9^th^ of February 2019 and were subjected to multiple sequence alignment using Bioedit for conservancy. The main analytical tool was IEDB, Chimera for homology modelling, and MOE for docking.

Among the tested peptides, fifteen promising T-cell peptides were predicted. Five peptides were more important than the others (HMIFASFIR, YRGVYPFIY, AVAAVSAAY, LRWAVSEAV, and IFASFIRTA) They show high Binding Affinity to MHC molecules, low binding energy required indicating more stable bonds, and their ideal length of nine peptides. (PTRAEVSDV) peptide is the most promising linear B-cell peptide due to its physiochemical parameters and optimal length (nine amino acids). It’s highly recommended to have these five strong candidates in future in vivo and in vitro analysis studies.

## 1. Introduction

The fungus *Candida albicans* is an opportunistic pathogen that causes a wide range of infections. *Candida albicans* is the primary cause of candidiasis and is the fourth most common cause of nosocomial infection. In addition, Disseminated invasive candidiasis that is a major complication of the disease has an estimated mortality rate of 40%-60% even with the use of antifungal drugs.^1–5^ it occurs with various other manifestations ranging from vaginal infections, which affect up to 75% of the women at least once in their lifetime, to deep systemic infections in hospitalized patients leading to high morbidity and mortality rates specially, in immunocompromised individuals, including organ recipients and HIV patients ^1, 2, 6–13^

Three proteins are most likely to be immunogenic in candida albicans which are the fructose-bisphosphate aldolase (Fba1), the phosphoglycerate kinase (Pgk) and the pyruvate kinase (Pk) with (pk) and (Fba1) being more immunodominant.^11, 14^ PK a 504 bp long, highly conservative protein which catalyzes the final step in glycolysis, converting phosphoenolpyruvate (PEP) and adenosine diphosphate (ADP) to pyruvate and adenosine triphosphate (ATP), once pyruvate is synthesized; it is either fed into the tri-carboxylic acid (TCA) cycle for oxidative phosphorylation or reduced to lactate under anaerobic conditions by lactate dehydrogenase enzyme LDH. ^15–17^

Vaccination is a significant approach to improve the standard of public health care and provide cost effective way to control the increasing numbers of infections. Many previous studies reached to the conclusion that C. albicans has become a multidrug resistant fungi. C. albicans growing in low oxygenated environments such as intravenous lines, renal catheters, bio prosthetic devices and self-organizing into biofilms which are resistant to a wide range of antifungal drugs, making conventional antifungal agents ineffective for the treatment of C. albicans infections.^1, 18–23^ Adding to that, the prolonged use of antifungal agents such as azoles, polyenes, and echinocandinsin in immuno-compromised individuals may lead to an increase in C. albicans resistance to many of the drugs used currently. ^4, 5, 11, 24^

Over the last decades, several different anti-Candida vaccines have been suggested with Different strategies for immunization against candidiasis such as, live-attenuated fungi, recombinant proteins, and glycoconjugates2526,27.Among many vaccines candidates, two univalent subunit vaccines (Als3 and the Sap2 v) and one recombinant C. albicans adhesin/invasin protein (NDV-3A) have been under development in recent years but none has been approved by the FDA, yet. The difficulties originates from the ability of C. albicans to evolve rapidly and lose host immune system’s recognition and the problem of how to trigger an immune responses in immunocompromised individuals. This calls for a new and a more effective version of these vaccines. ^24, 27–32^

This study aims to introduce a new possible vaccine for *C. albicans* through analyzing peptides of its pyruvate kinase (PK) protein as an immunogenic stimulant computationally. Vaccine production that depends on biochemical experiments can be expensive, time consuming, hazardous and more likely to give unsatisfactory results, that’s why bioinformatics approaches provide more affordable, efficient and faster techniques for vaccine development.^33–35^ Using immunoinformatics approaches, otherwise known as computational immunology which is an emerging specialization of bioinformatics that has recently been used to predict possible peptide based vaccines using massive online databases. This study is different than other studies for being the first in scilico analysis study of its kind attempting to design a peptide-based vaccine for *C. albicans* using Pyruvate kinase (PK) as an immunogenic stimulant.

## METHODOLOGY

### 1.1. Sequence retrieval

A total number of 28 *candida albicans*, pyruvate kinase proteins were obtained from national center of biotechnology information NCBI on the 9^th^ of February 2019. (Available at: https://www.ncbi.nlm.nih.gov/)

### 1.2. Multiple sequence Alignment

A multiple sequence alignment was made using the Clustal w package, Bioedit tool for the sequences by blasting them against reference sequence (accession: XP_714934.1) to show areas of protein conservation in the different species and exclude peptides located in areas of low conservation. Peptides located at highly conserved regions are more likely to give stronger vaccine that covers more populations than peptides at low areas of conservation.^36, 37^

### 1.3. B-cell epitopes prediction

#### 1.3.1. Linear epitopes

Was made through Immune Epitope Database Analysis Resource IEDB tool for Linear b-cell epitopes prediction. The tool has different methods to predict epitopes antigenicity, flexibility, immunogenicity, and surface accessibility. This this done through the submission of the Swiss-prot ID of the reference protein or FASTA. Then peptides larger than six peptides were spliced to increase the possibility of obtaining peptides with higher scores in Bepipred, Kolaskar & Tongaonka, and Emini. Finally, a total number of 144 peptides were obtained, 129 of which were conserved and were put to the Bepipred surface test and fifteen peptides were located in non-conserved regions and sequentially were removed. (Available at: http://tools.iedb.org/main/bcell/)

##### 1.3.1.1 Bepipred linear epitope prediction

Peptides were introduced to this test to check their linearity, linear peptides are simple, unbranched, where non-linear peptides are found in clusters. Linear peptides that are located on surface have higher probability of provoking the immune system to induce a response than non-linear or peptides which are not on surface. Peptides that passed the linearity test scored above the threshold of one, were 106 peptides, while the 23 other peptides were excluded for scoring less than one.^38–40^

##### 1.3.1.2. Emini surface accessibility test

It’s a test that calculates surface probability using a specific formula. It was performed to linear peptides composed of six amino acids and had a threshold of one, 81 peptides passed while 25 peptides were not on the surface and were not further analyzed.^37^

##### 1.3.1.2 Kolaskar & Tongaonka Antegenicity test

It’s a test that uses physiochemical properties of amino acids residues and the frequency of their appearance in peptides under study to predict protein antigenicity. The 81 peptides were submitted to the Antigenicity test with a threshold of 1.033, and five peptides passed indicating they are capable of inducing an immune response. ^41^

#### 1.3.2. Non-linear B-cell epitopes

##### 1.3.2.1 ElliPro Antibody epitopes prediction

It’s a tool that works using the PDB ID of the protein, providing minimum score of (0.5) and maximum distance of (6) to predict non-linear peptides. It provides 3D model of the clustered peptides with the result. ^42^ (available at: http://tools.iedb.org/ellipro/)

#### 1.4 T-cell epitopes prediction

##### 1.4.2 Binding to MHC class I prediction tool

Is a tool which predicts peptides binding to MHC-I molecules following the submission of sequence, prediction method (Artificial neural network ANN method was used in this study), source of MHC molecules (Homo sapiens) and frequently occurring alleles, and HLA sets of twenty-seven alleles that are desired in the study. The result is then sorted by percentile rank and cutoff below IC50 of 500.^43–48^ (Available at: http://tools.iedb.org/mhci/)

##### 1.4.3 Binding to MHC class II prediction tool

It’s a similar tool to MHC I tool, after submitting the sequence, prediction method (NN-align algorithm used), selecting the source and locus, and MHC alleles. The result is then sorted by percentile rank and a cutoff of IC50 of 100.^49^ (Available at: http://tools.iedb.org/mhcii/)

##### 1.4.4 Population coverage

T-cell recognizes a complex of major histocompatibility complex (MHC) molecules. This tool is used to test the ability of a peptide to elicit a response in immune system in specific populations carrying the desired MHC allele or worldwide. This tool calculates the approximate percentage of individuals’ response to an epitope with regard to ethnicity and HLA genotypic frequencies and T-cell restriction. Population coverage is tested throughout the whole protein to MHC molecules class I, II and combined binding.^50^ (Available at: http://tools.iedb.org/population/)

### 2.5. Molecular docking analysis

Molecular docking was performed using Moe 2007 Accessed July 22, 2019. The 3D structures of the promiscuous epitopes were predicted by PEP-FOLD.^51, 52^ The crystal structure of HLA-A*68:01 (**PDB ID 4hwz**) and HLA-DRB1*01:01(**PDB ID 5jlz**) were chosen as a model for molecular docking and were downloaded in a PDB format from the RCSB PDB resource. However, the selected crystal structures were in a complex form with ligands. Thus, to simplify the complex structure all water molecules, hetero groups and ligands were removed by Discovery Studio Visualizer 2.5.^53^ Partial charge and energy minimization were applied for ligands and targets. In terms of the identification of the binding groove, the potential binding sites in the crystal structure were recognized using the Alpha Site Finder. Finally, ten independent docking runs were carried out for each Peptide. The results were retrieved as binding energies. Best poses for each epitope that displayed lowest binding energies were visualized using UCSF chimera 1.13.1 software.^54^ (available at: https://www.chemcomp.com/, and http://bioserv.rpbs.univ-paris-diderot.fr/services/PEP-FOLD3/)

### 2.6. Physiochemical parameters

Hence, the main function of a vaccine is to induce an immunogenic response once introduced to the immune system, it is essential to recognize the physiochemical parameters of the protein it’s made of. This has been done using protoparam which has been used to give information about the vaccine of PK (molecular weight, theoretical pI, amino acid composition, estimated half-life, instability index, aliphatic index and grand average of hydropathicity (GRAVY)), Protscale which provided hydrophobicity information and graphs (Both softwares are found in ExPasy server) And Bioedit software that provided the amino acid composition graph of PK protein ^36, 37, 55^ (Available at: https://web.expasy.org/protparam/, and https://web.expasy.org/protscale/)

### 2.7 Homology modelling

Using **Chimera** (version 1.13.1rc) a tool powered by University of California, San Francisco, with support from NIH P41-GM103311. Chimera is used to visualize the 3-dimentional structure of the pyruvate kinase (PK) protein of C. albicans and the most promising peptides binding to MHC class I, MHC class II and peptides binding to MHC class I and II altogether in the study. (Figures 11,12 and 13)^56,57,58^

## 3. Results

### 3.1. B-cell epitope prediction

The reference sequence of Pyruvate kinase *C. albicans* was analyzed using Bepipred Linear Epitope Prediction first, the average binder’s score of the protein to B cell was 0.076, minimum was −0.002 and 1.916 for a maximum score, all values equal or greater than the default threshold 0.350 were potentially linear epitopes (table 1 and figure 2).

**Figure 1:**
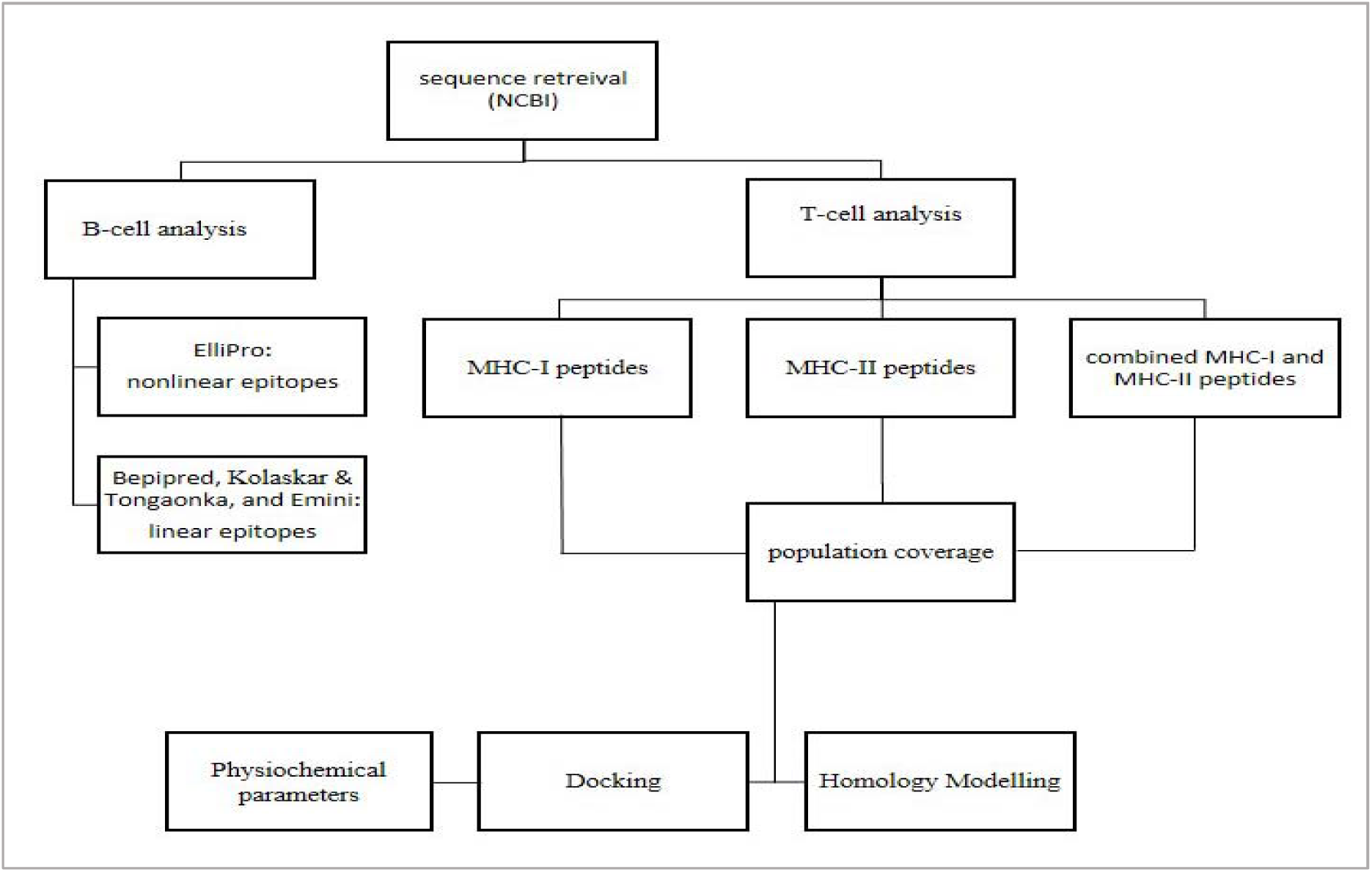
a hierarchy template to visualize the work flow followed in the study

**Figure 2:**
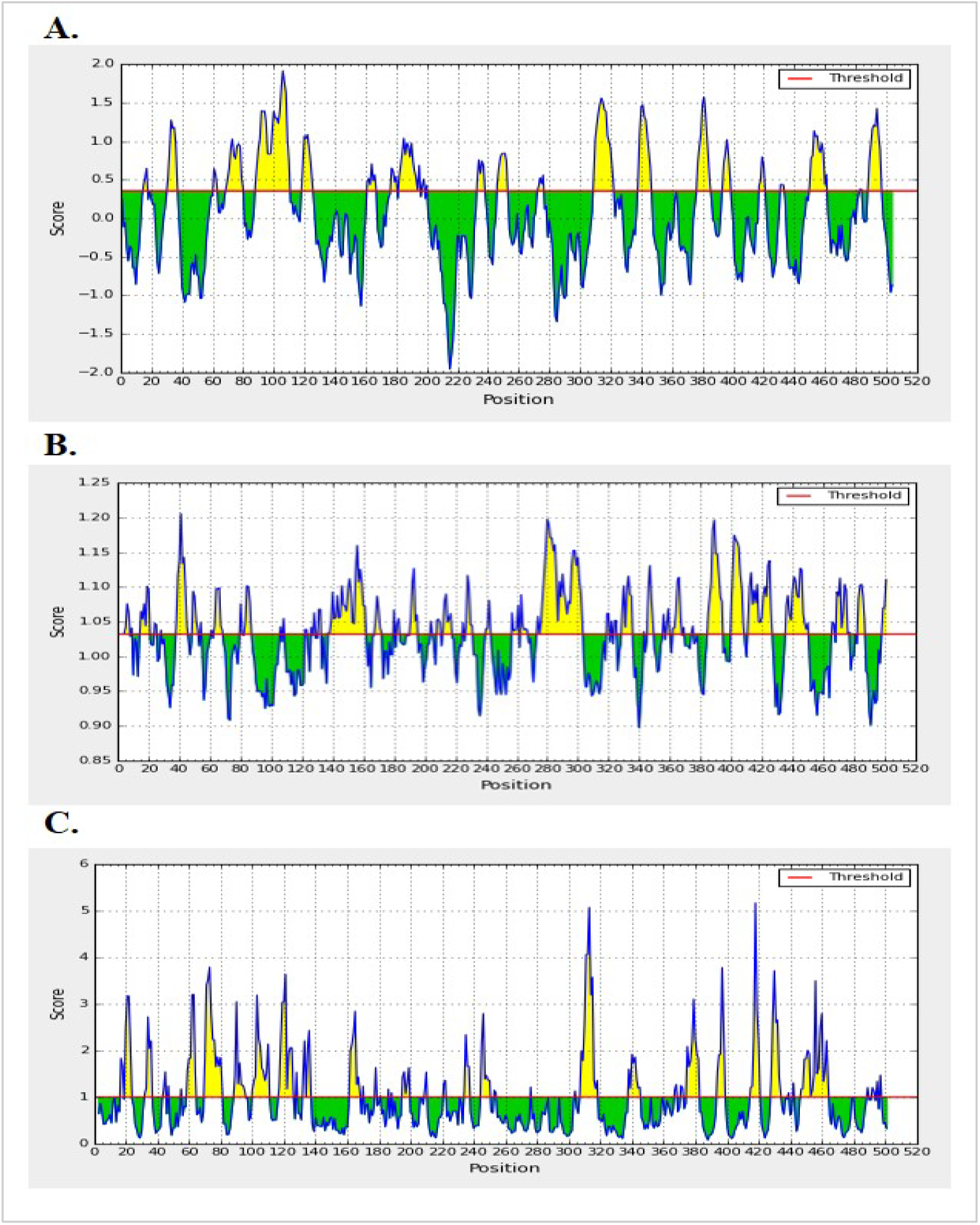
**A.** Bepipred Linear Epitope Prediction of PK *C. albicans*, the x-axis and y-axis represent the sequence position and linear probability, respectively. **B.** EminI surface accessibility prediction of PK *C. albicans*, the x-axis and y-axis represent the sequence position and surface probability, respectively. **C.** Kolashkar and Tongaonkar antigenicity prediction of PK Candida albicans, the x-axis and y-axis represent the sequence position and antigenic propensity, respectively. Yellow peaks above the threshold (Red line) passed the test, Green peaks fall below threshold and did not pass the test.

**Table 1:** Predicted antigenic B-cell epitopes, 12 antigenic sites were identified from PK *C. albicans*

| Start | End | Peptide | Length |
| --- | --- | --- | --- |
| 161 | 166 | VDDEQT | 6 |
| 393 | 398 | AAYEQD | 6 |
| 31 | 37 | IGPKTNN | 7 |
| 119 | 125 | DAYKTKC | 7 |
| 246 | 252 | ENQQGVN | 7 |
| 338 | 346 | ETAKGNYPV | 9 |
| 377 | 385 | KKPTATTET | 9 |
| 488 | 497 | GWTRGSGHSN | 10 |
| 450 | 461 | KPSIENWQEDVE | 12 |
| 182 | 195 | GVNLPGTDVDLPAL | 14 |
| 309 | 322 | TYNPRPTRAEVSDV | 14 |
| 88 | 110 | DTKGPEIRTGTTIGDKDYPIPPN | 23 |

In Emini surface accessibility prediction, for a potent B-cell epitope the average surface accessibility areas of the protein was scored as 1.000, with a maximum of 5.174 and a minimum of 0.073, all values equal or greater than the default threshold 1.000 were potentially in the surface. In addition, Kolaskar and Tongaonkar antigenicity prediction’s average of antigenicity was 1.033, with a maximum of 1.206 and minimum of 0.897; all values equal to or greater than the default threshold 1.033 are potential antigenic determinants. The results of all conserved predicted B cell epitopes are shown in (table 2 and figure 2).

**Table 2:**
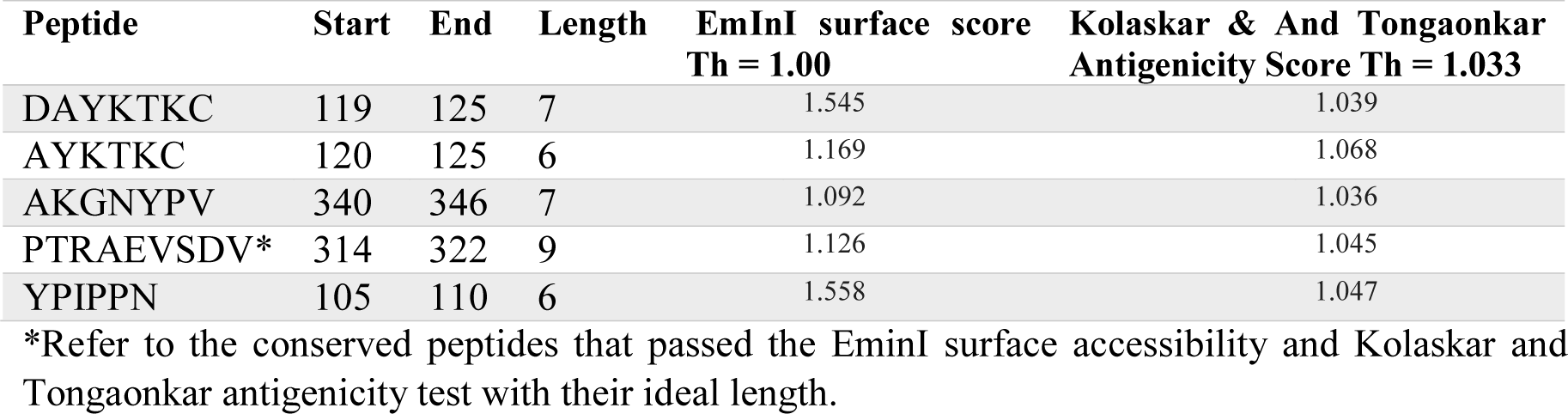
B-cell epitopes prediction with their surface accessibility score and antigenicity score:

Though five linear conserved surface antigenic epitopes passed all of the above tests, one epitope out of all was thought to be the promising B-cell epitope that is able to evoke B-lymphocyte for their efficient physiochemical properties and length (PTRAEVSDV). In addition, eighteen promising discontinuous epitopes were defined from the modeled protein after submission to ElliPro prediction tool. Epitopes were predicted to be located on the surface of the protein indicating quick recognition by host immune system. (Table 3, figure 3)

**Figure 3:**
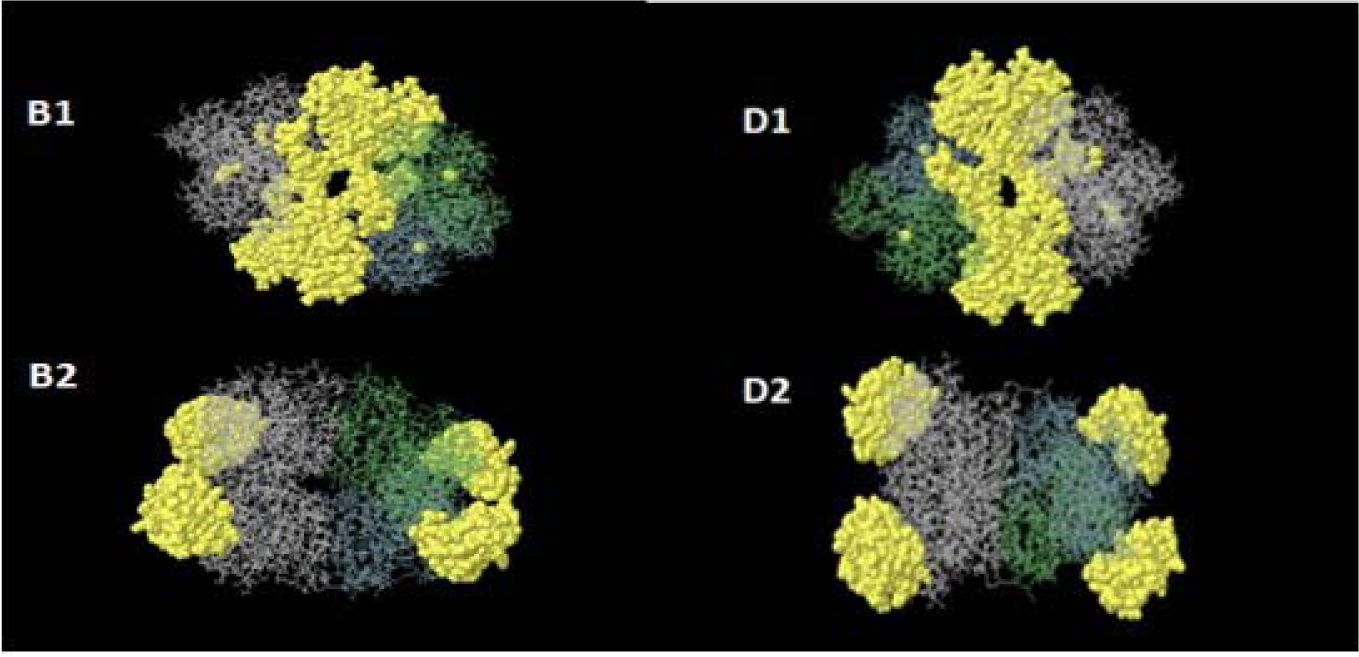
Three-dimensional representation of four non-linear epitopes (B1–B2, D1-D2) of the highly immunogenic pyruvate Kinase PK protein of *C. albicans* using ElliPro prediction tool. The clustered epitopes are highlighted in yellow, and the bulk of PK protein is depicted in grey.

**Table 3:**
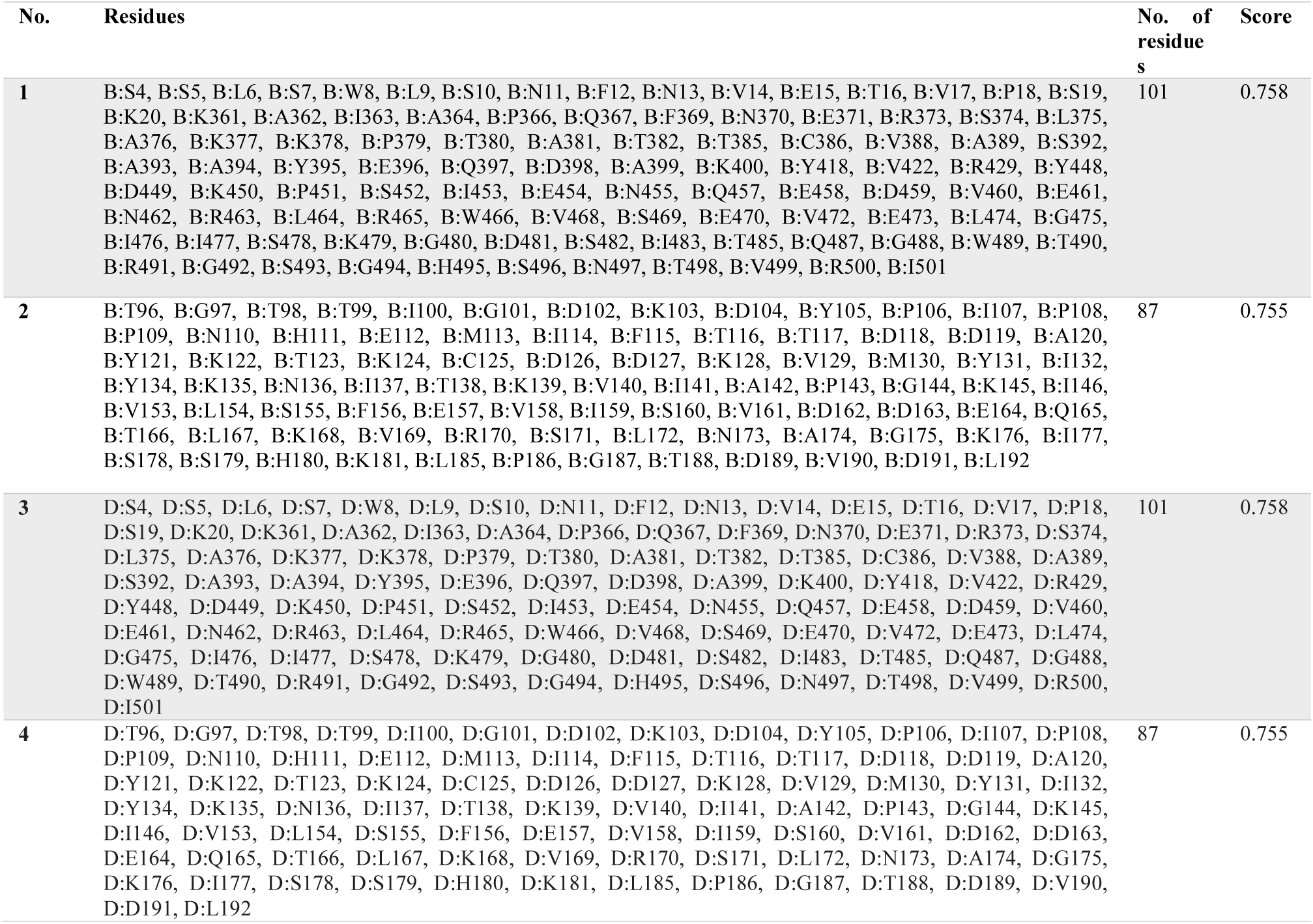
List of four predicted non-linear B-cell epitopes with highest number of Residues and their scores using ElliPro tool

### 3.2. T-cell peptide predictions

#### 3.2.1. Prediction of MHC-I binding profile for T cytotoxic cell conserved epitopes

167 epitopes were anticipated to interact with di erent MHC-1 alleles. The core epitopes (HLYRGVYPF/ RAAKFSHLY) were noticed to be the dominant binders with 10 alleles for each (HLA-A*02:06,HLA-A*23:01, HLA-A*29:02,HLA-A*32:01, HLA-B*15:01,HLA-B*15:02, HLA-B*35:01, HLA-C*07:02, HLA-C*12:03, HLA-C*14:02, HLA-A*11:01, HLA-A*29:02, HLA-A*30:02, HLA-A*68:01, HLA-B*15:01, HLA-B*35:01, HLA-B*57:01, HLA-B*58:01, HLA-C*12:03, HLA-C*15:02), followed by YVDDGVLSF which binds with nine alleles, AVAAVSAAY binding with eight alleles and YRGVYPFIY that’s believed to bind with five alleles. (Table 4).

**Table 4:**
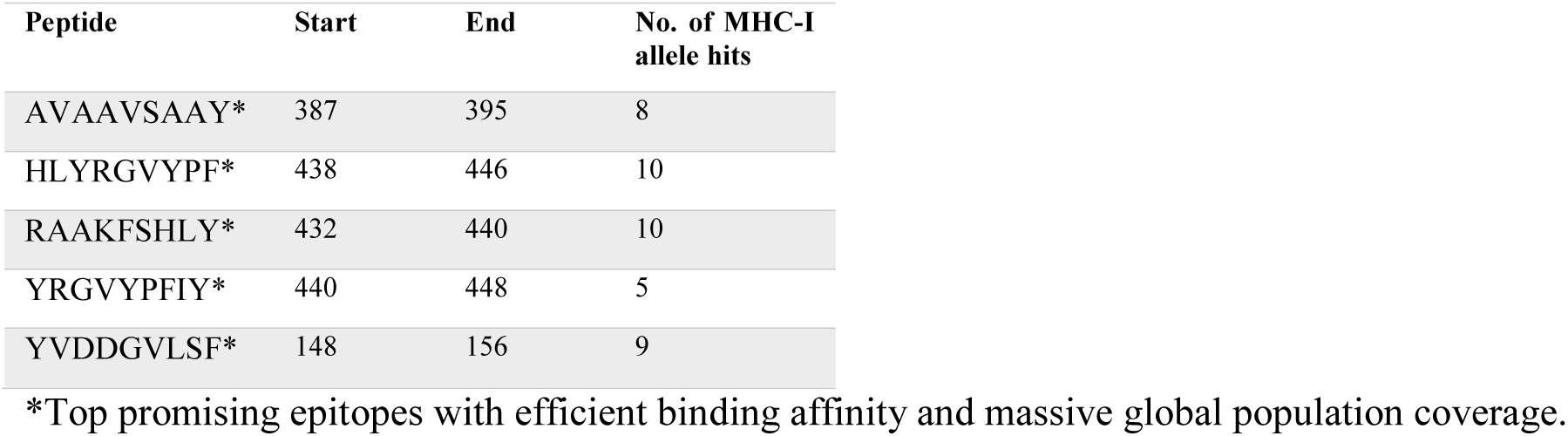
List of most promising epitopes that had a good binding affinity with MHC-I alleles in terms of IC50 and Percentile rank.

| Peptide | Start | End | No. of MHC-I allele hits |
| --- | --- | --- | --- |
| AVAAVSAAY* | 387 | 395 | 8 |
| HLYRGVYPF* | 438 | 446 | 10 |
| RAAKFSHLY* | 432 | 440 | 10 |
| YRGVYPFIY* | 440 | 448 | 5 |
| YVDDGVLSF* | 148 | 156 | 9 |

\*Top promising epitopes with efficient binding affinity and massive global population coverage.

#### 3.2.2. Prediction of MHC-II binding profile for T helper cell conserved epitopes

125 conserved predicted epitopes were found to interact with MHC-II alleles. The core epitopes (HMIFASFIR) is thought to be the top binder as it interacts with nine alleles; (HLA-DPA1*01, HLA-DPB1*04:01, HLA-DPA1*01:03, HLA-DPB1*02:01, HLA-DPA1*02:01, HLA-DPB1*01:01, HLA-DPA1*02:01, HLA-DPB1*05:01, HLA-DRB5*01:01). Followed by LRWAVSEAV which binds to seven alleles and (IAYPQLFNE, IFASFIRTA, VFVVQKQLI) binding to six alleles for each. (Table 5).

**Table 5:** List of the five promising core sequence epitopes that had a strong binding affinity with MHC-II in terms of IC50 and Percentile Ranks

| peptide Sequence | No. of MHC-II allele hits |
| --- | --- |
| HMIFASFIR | 9 |
| IAYPQLFNE | 6 |
| IFASFIRTA | 6 |
| VFVVQKQLI | 6 |
| LRWAVSEAV | 7 |

#### 3.2.3. Molecular docking analysis

T-cell epitopes were tested to visualize the binding affinity between promising epitopes and HLA molecules. Global energy is the energy required to estimate the strength of association between the epitope within the active cleft of MHC molecule; more negative value indicates favored and stable binding of the complex. (Table 6, Figure 4, and 5)

**Figure 4.**
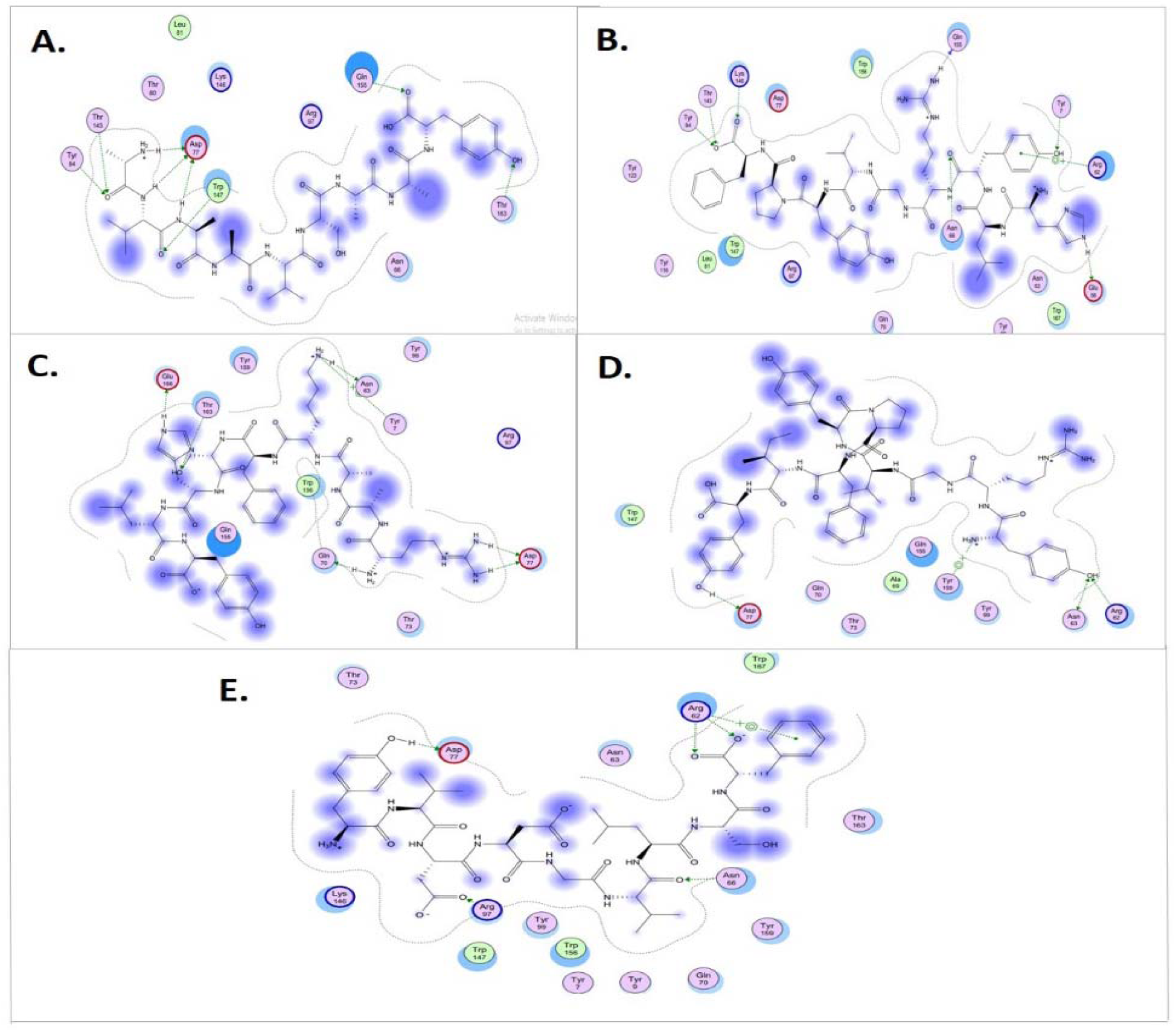
Illustrates 2-dimentional interaction of the best docking poses with HLA-A*68:01 MHC-I allele for five promising peptides **A.** AVAAVSAAY **B.** HLYRGVYPF **C.** RAAKFSHLY **D.** YRGVYPFIY **and E.** YVDDGVLSF.

**Table 6.**
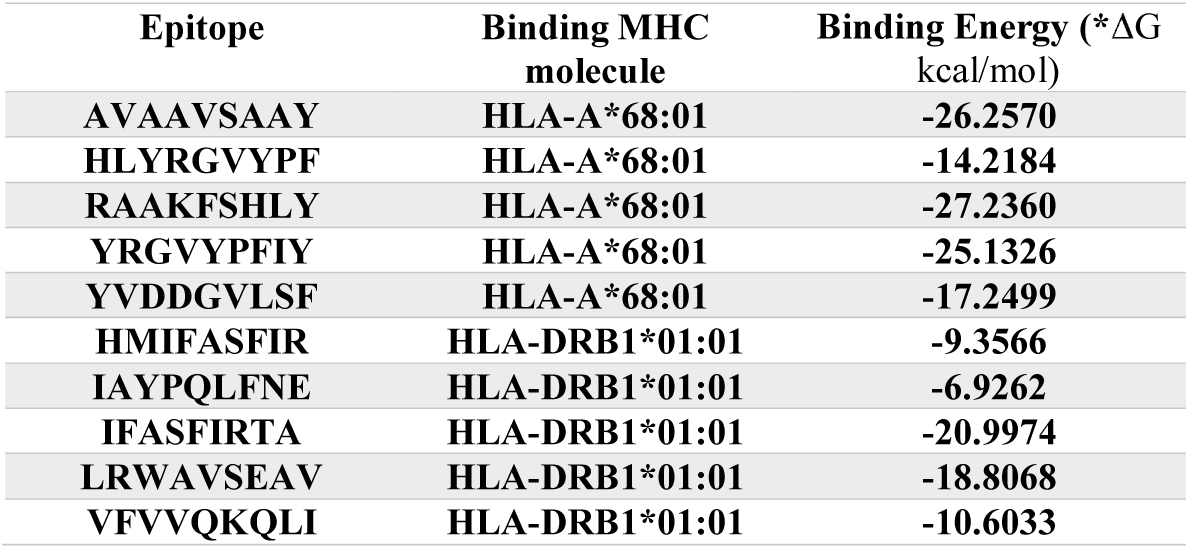
Docking results of the most promiscuous epitopes that show the best binding affinity

### 3.4. Physiochemical parameters

The length of the protein was found to be 504 amino acids. Molecular weight and Theoretical pI parameters were calculated as 55071.50KDa and 7.17, respectively. The pI value reflects that the protein nature is acidic. The total numbers of negatively and positively charged residues estimated as 65 and 65, correspondingly. Extinction coefficient of protein at 280 nm was measured 25035 M-1 cm-1 in water. The estimated half-life of vaccine was; 30 hours in (mammalian reticulocytes, in vitro),>20 hours in (yeast, in vivo) and >10 hours in (Escherichia coli, in vivo). The Instability index was computed to be 27.29, which indicates the thermo-stability of protein. The aliphatic index and the grand average of hydropathicity (GRAVY) value of vaccine were determined 91.19 and (−0.151), respectively. The high aliphatic index shows that the protein is stable in a wide range of temperatures, and the negative GRAVY value shows protein hydrophilicity and a better interaction with the surrounding water molecules. (Illustrated in table 7, figure 5, 6).

**Figure 5.**
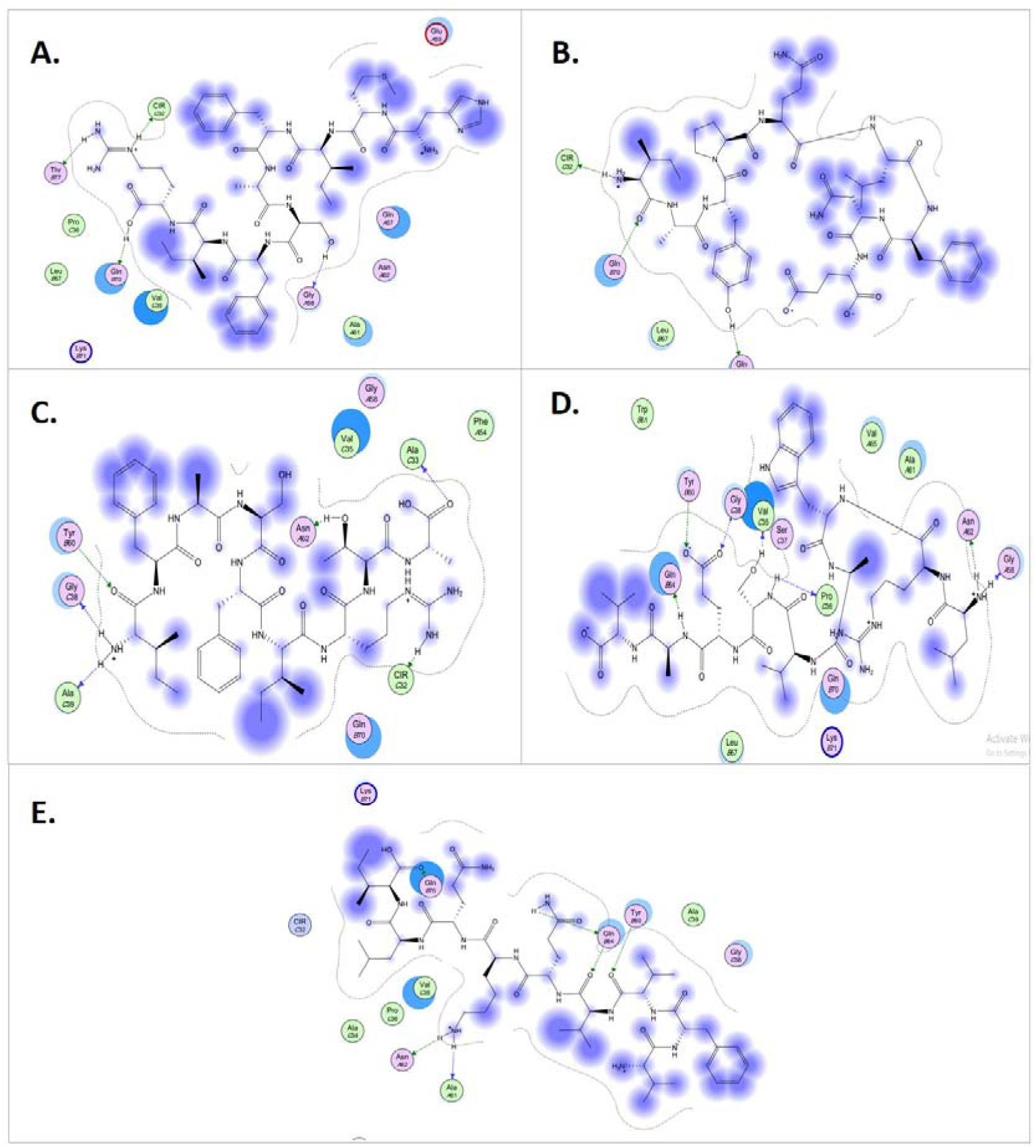
Illustrates 2-dimentional interaction of the best docking poses with HLA-DRB1*01:01 MHC-II allele for five promising peptides **A.** HMIFASFIR **B.** IAYPQLFNE **C.** IFASFIRTA **D.** LRWAVSEAV **and E.** VFVVQKQLI

**Figure 6.**
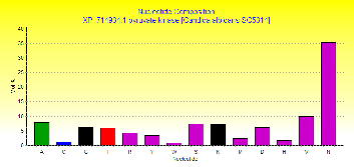
A graph illustrating the Amino Acid Composition (Mol %) of PK protein using BioEdit sequence alignment tool (Version 7.2.5.)

**Table 7.** List of amino acid that formed pyruvate kinase protein, their sequential location, and molar percentage (Mol %)

| AA | Number | Mol% | AA | Number | Mol% |
| --- | --- | --- | --- | --- | --- |
| **Ala (A)** | 54 | 10.7 | **Leu (L)** | 37 | 7.3 |
| **Arg (R)** | 29 | 5.8 | **Lys (K)** | 36 | 7.1 |
| **Asn (N)** | 16 | 3.2 | **Met (M)** | 15 | 3 |
| **Asp (D)** | 28 | 5.6 | **Phe (F)** | 16 | 3.2 |
| **Cys (C)** | 10 | 2 | **Pro (P)** | 21 | 4.2 |
| **Gln (Q)** | 15 | 3 | **Ser (S)** | 27 | 5.4 |
| **Glu (E)** | 37 | 7.3 | **Thr (T)** | 26 | 5.2 |
| **Gly (G)** | 37 | 7.3 | **Trp (W)** | 2 | 0.4 |
| **His (H)** | 12 | 2.4 | **Tyr (Y)** | 9 | 1.8 |
| **Ile (I)** | 38 | 7.5 | **Val (V)** | 39 | 7.7 |

### 3.5. Population coverage analysis:-

The most interesting findings in this test is that the population coverage analysis result for most common binders to MHC-I and MHC-II alleles per each and combined; exhibit an exceptional coverage with percentages of 93.11%, 99.7%, and 99.7%. Respectively.

#### 3.5.1. Population coverage for isolated MHC-I

Five epitopes are given to interact with the most frequent MHC class I alleles (HLYRGVYPF, RAAKFSHLY, YVDDGVLSF, AVAAVSAAY and YRGVYPFIY), representing a considerable coverage against the whole world population. The maximum population coverage percentage over these epitopes worldwide was found to be 54.79% for YRGVYPFIY (table 8, figure 7).

**Figure 7.**
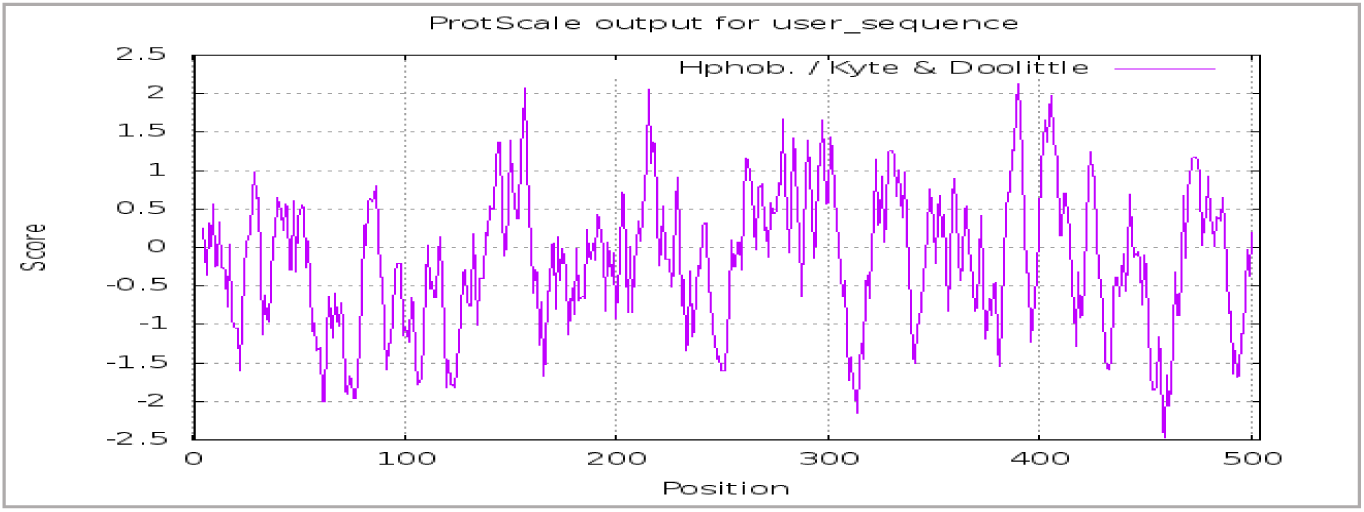
A graph showing the Hydrophobicity scale of PK protein using ProtScale server.

**Table 8.**
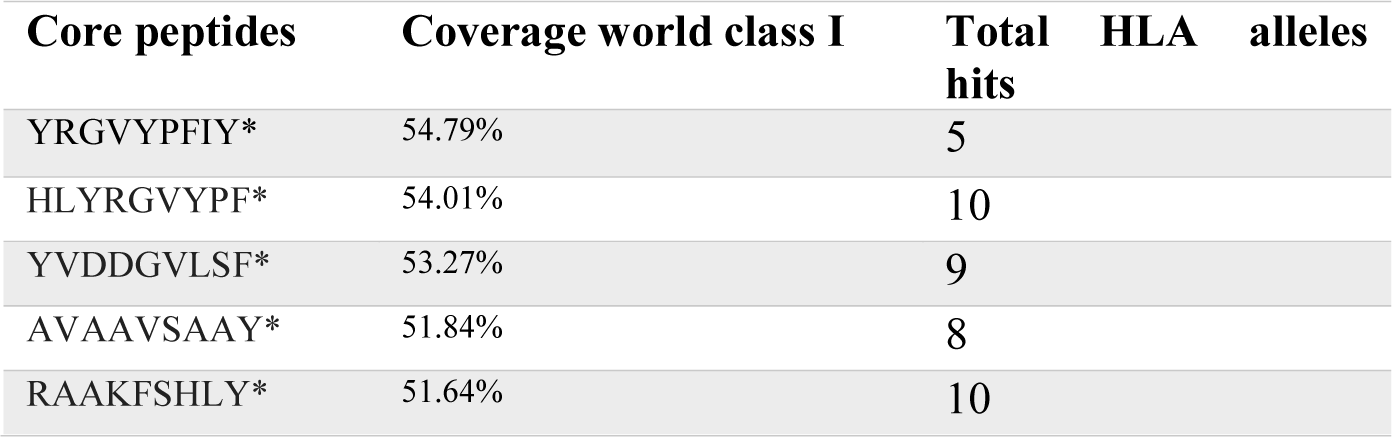
Global Population coverage of promising epitopes binding to MHC class I alleles

#### 3.5.2. Population coverage for isolated MHC-II

In the case of MHC class II, five epitopes were assumed to interact with the most frequent MHC class II alleles (HMIFASFIR, LRWAVSEAV, IAYPQLFNE, IFASFIRTA and VFVVQKQLI), inferring a massive global coverage. The highest population coverage percentage of these epitopes worldwide was that of HMIFASFIR with percentage of 95.03% (table 9, figure 8).

**Figure 8.**
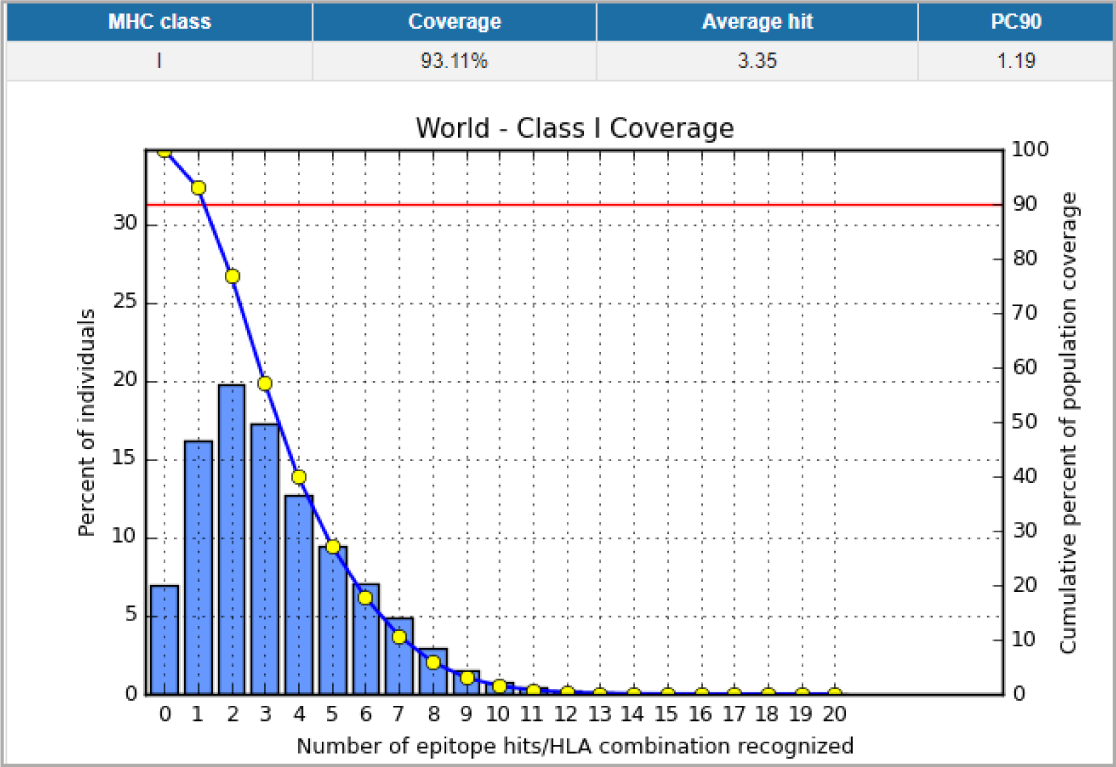
Illustrates the global coverage for the top five MHC-I peptides (HLYRGVYPF, RAAKFSHLY, YVDDGVLSF, AVAAVSAAY and YRGVYPFIY). Note: In the graph, the line (-o-) represents the cumulative percentage of population coverage of the epitopes; the bars represent the population coverage for each epitope.

**Table 9.**
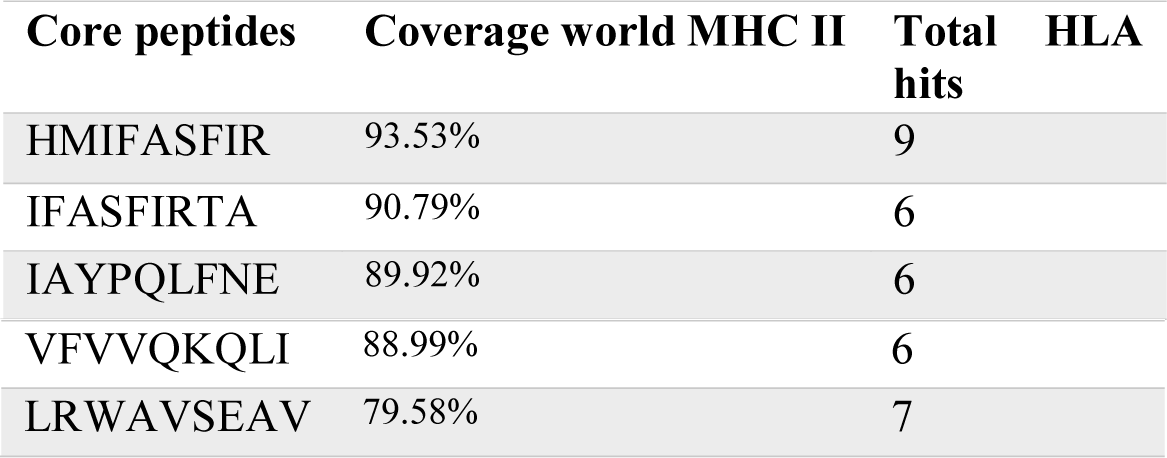
Global Population coverage of promising epitopes in isolated MHC class II

#### 3.5.3. Population coverage for MHC-I and MHC-II alleles combined

Regarding MHC-I and MHC-II alleles combined, three epitopes were supposed to interact with most predominant MHC class I and MHC class II alleles (HMIFASFIR, TETCAVAAV and LRWAVSEAV); represent a significant global coverage by IEDB population coverage tool. The most common population coverage percentage of these epitopes in the World was granted to HMIFASFIR with percentage of 97.02%. The remarkable finding in this test is the average for the most common binders to combined MHC-I and MHC-II alleles; reveal an outstanding coverage with percentage of 99.75%.Table 10, Figure 9

**Figure 9.**
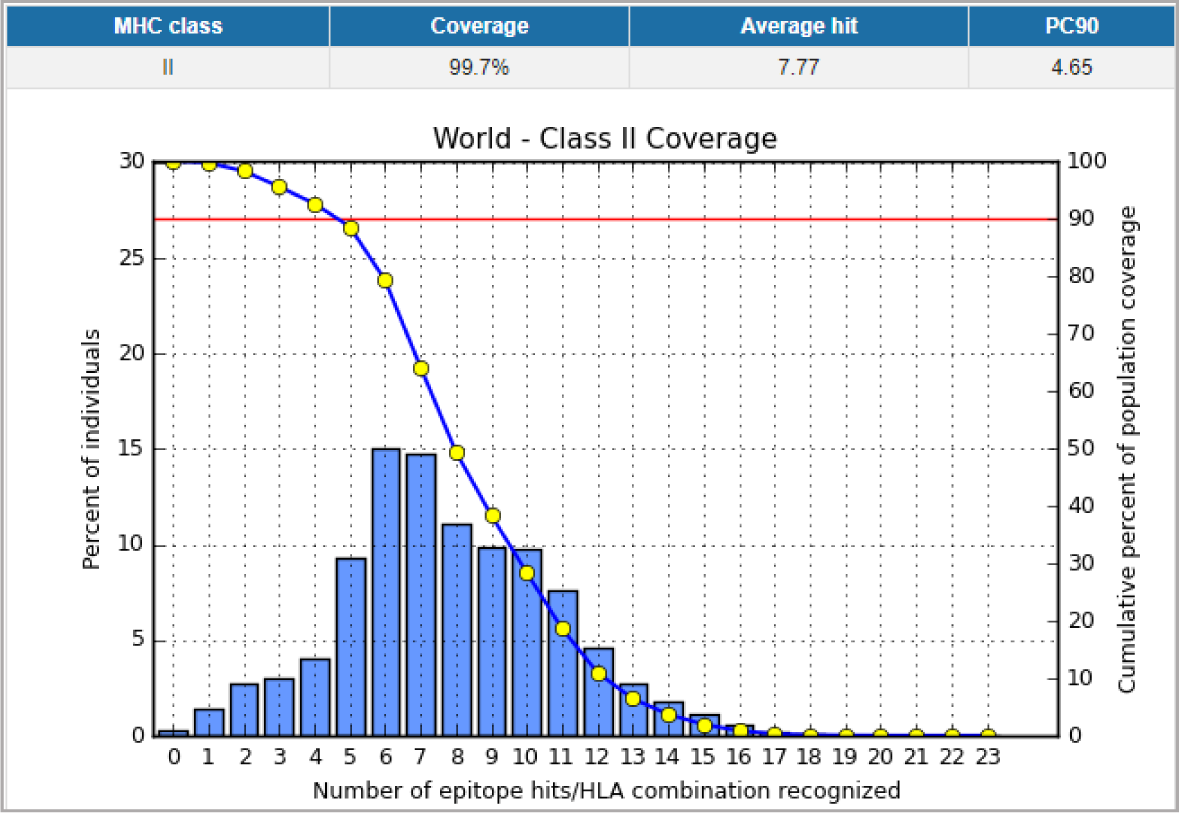
Illustrates the global proportion for the top five MHC-II epitopes (HMIFASFIR, LRWAVSEAV, IAYPQLFNE, IFASFIRTA and VFVVQKQLI). Notes: In the graph, the line (-o-) represents the cumulative percentage of population coverage of the epitopes; the bars represent the population coverage for each epitope.

**Figure 10.**
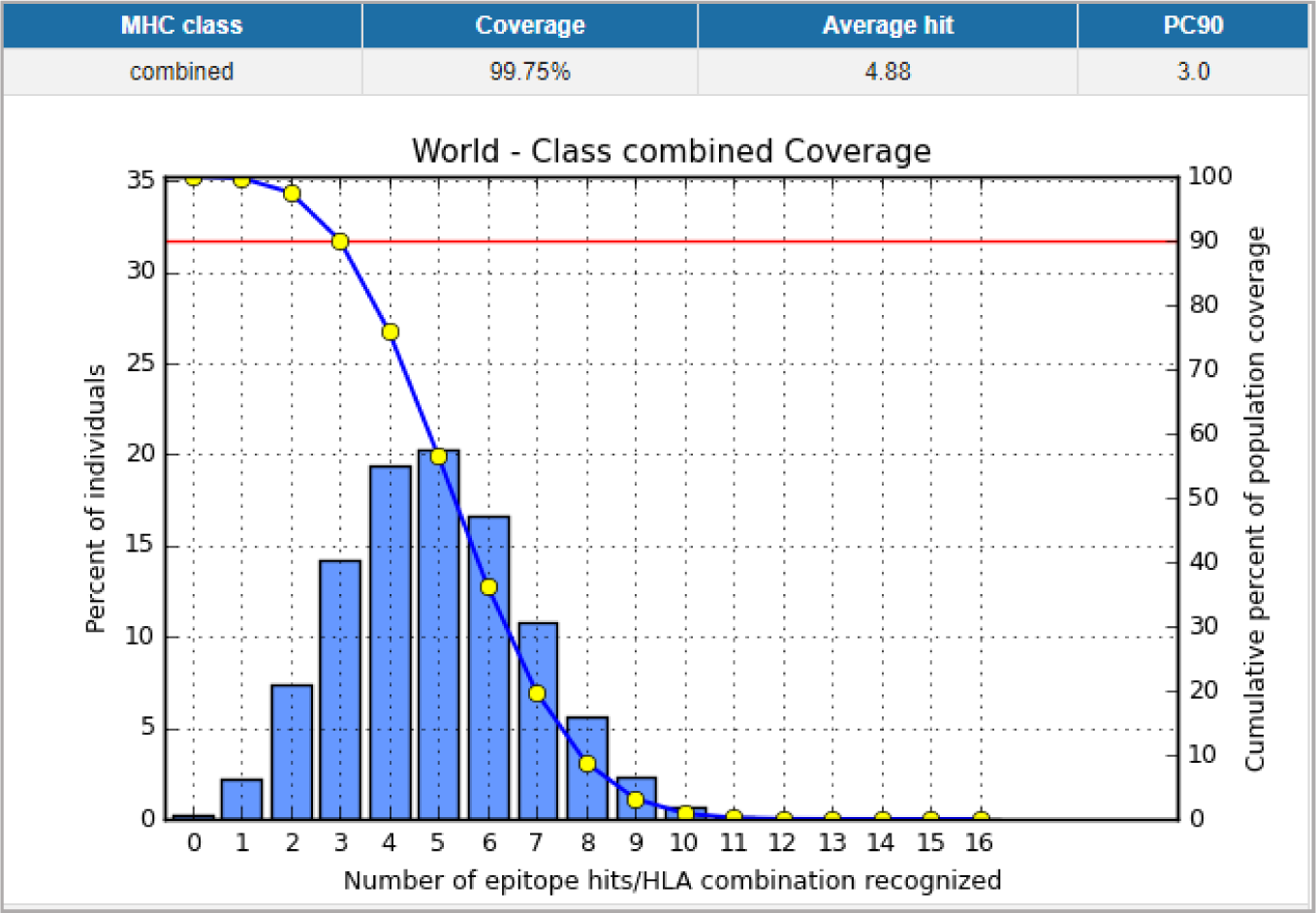
Illustrates the global population proportion for the top three MHC-I & II epitopes in combined mode (HMIFASFIR, TETCAVAAV and LRWAVSEAV). Notes: In the graphs, the line (-o-) represents the cumulative percentage of population coverage of the epitopes; the bars represent the population coverage for each epitope.

**Figure 11.**
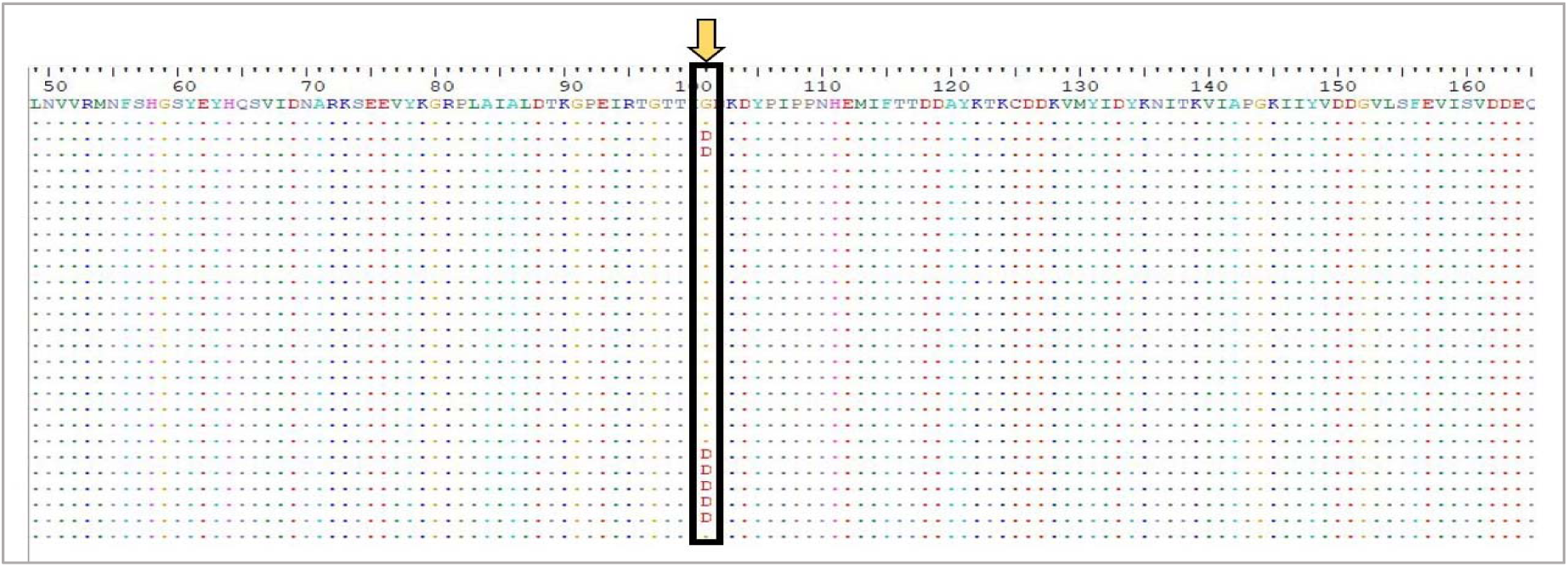
A fragment of the multiple sequence alignment of pyruvate kinase protein of *C. albicans* showing the only area of non-conservation in some species at position 101, where Glycine mutated into Aspartic acid, using Bioedit software.

**Figure 12.**
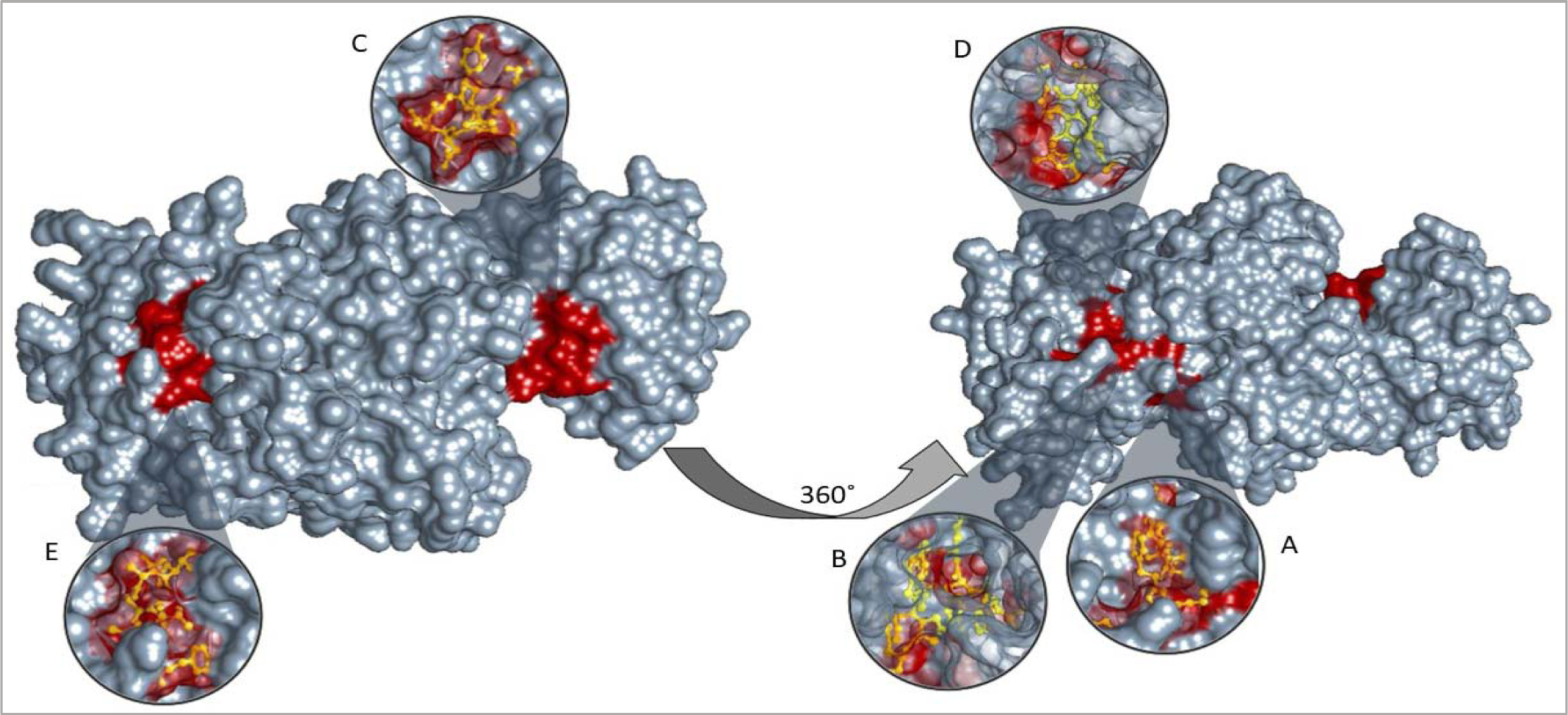
3D structure of pyruvate kinase protein visualizing top five T-cell peptides binding to MHC class I **A.** YRGVYPFIY **B.** HLYRGVYPF **C.** YVDDGVLSF **D.** RAAKFSHLY and **E.** AVAAVSAAY using chimera (version 1.13.1rc)

**Figure 13.**
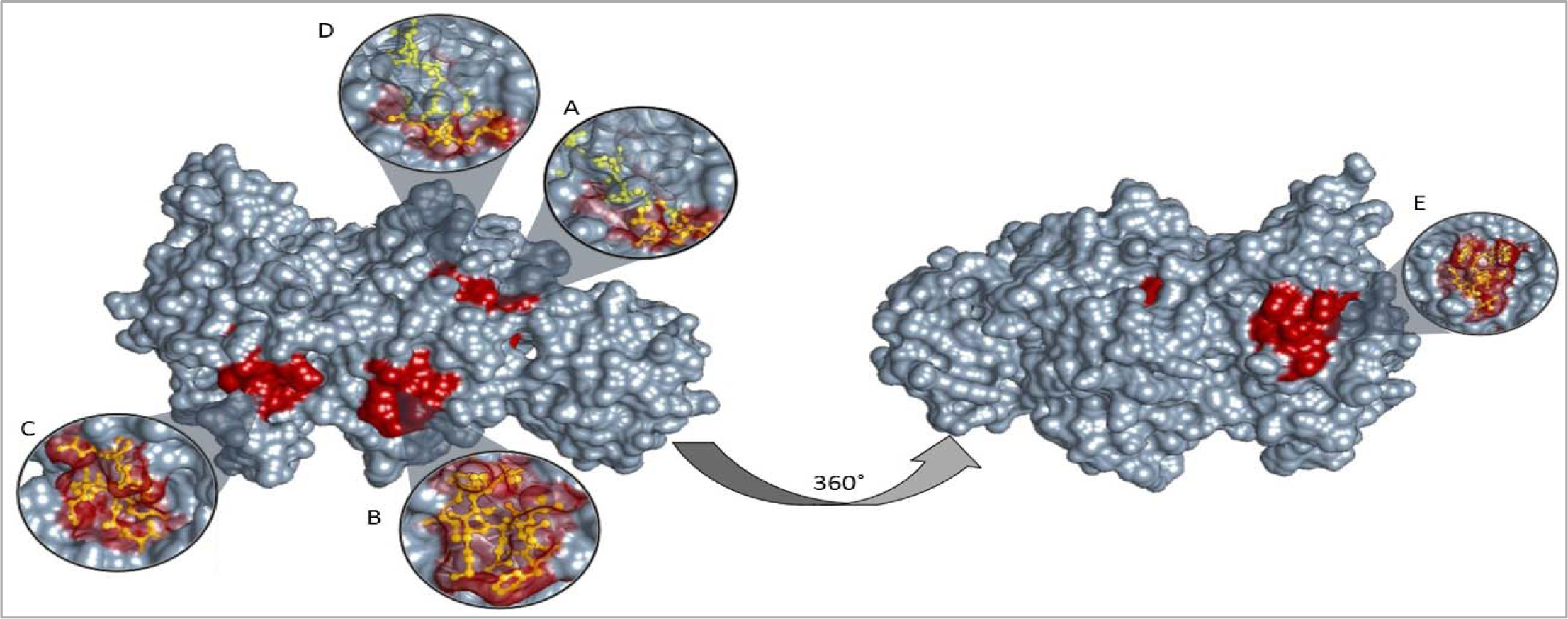
3D structure of pyruvate kinase protein visualizing top five T-cell peptides binding to MHC class II **A.** HMIFASFIR **B.** VFVVQKQLI **C.** IAYPQLFNE **D.** IFASFIRTA **and E.** LRWAVSEAV, using chimera (version 1.13.1rc)

**Figure 14.**
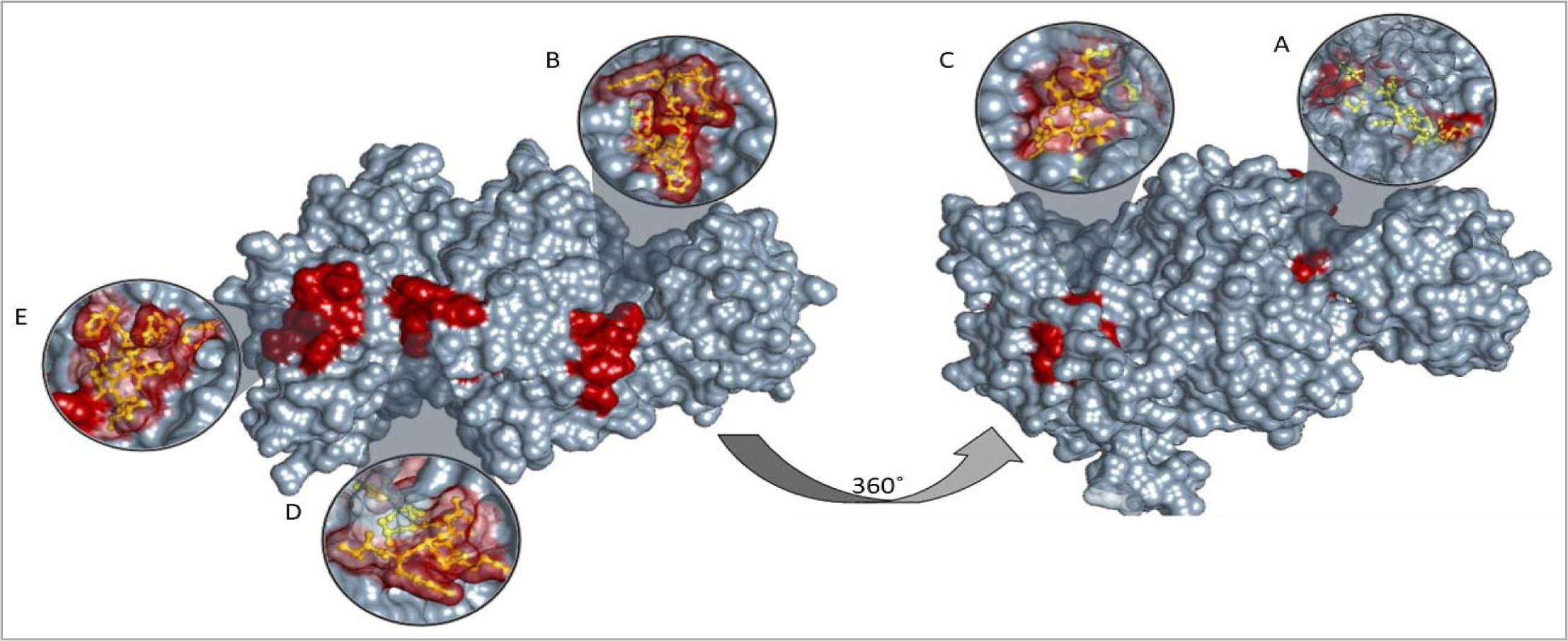
3D structure of pyruvate kinase protein visualizing top five T-cell peptides binding to MHC class I and II altogether **A.** HMIFASFIR **B.** NFSHGSYEY **C.** TETCAVAAV **D.** VYKGRPLAI **and E.** LRWAVSEAV, using chimera (version 1.13.1rc)

**Table 10.** Global population coverage of the ten promising epitopes in MHC class I and II combined mode.

| Core peptides | Global Coverage MHC classes I-II Combined | Total HLA hits |
| --- | --- | --- |
| HMIFASFIR | 97.02% | 13 |
| TETCAVA AV | 80.74% | 8 |
| LRWAVSEAV | 80.56% | 8 |

## 4. Discussion

This study with a major focus on T-cell epitopes has revealed fifteen promising T-cell peptides to make a peptide based vaccine for *C. albicans* using pyruvate kinase as a target protein to induce an immune response. The peptide HMIFASFIR is one of the most important findings of this study regarding its interaction with both MHC-II alleles and MHC-I and MHC-II alleles combined. Giving significantly high worldwide population coverage percentages of 95.03% and 97.03% respectively, ideal length of nine amino acids, an IC50 of 8.2 when binding to MHC allele HLA-DRB5*01:01 indicating strong interaction between the two and a binding energy of −9.356 (kcal/mol) when binding to HLA DRB1*01:01. In addition, We suggest PTRAEVSDV peptide is the most promising linear B-cell peptide due to its physiochemical parameters and optimal length (nine amino acids).

Five T-cell peptides were suggested to be the most promising ones in accordance to their global population coverage, binding energies to most frequent MHC-I allele in the study (HLA A*68:01) and their strength of interaction to other alleles with the IC50 scores. These epitopes could potentially induce CD8+ cytotoxic T-cell immune response when interacting strongly with MHC class I alleles. YRGVYPFIY, HLYRGVYPF, YVDDGVLSF, RAAKFSHLY, and AVAAVSAAY (having global population coverage of 54.79%, 54.01%, 53.27%, 51.84%, 41.64%, respectively) all five peptides had a massive global coverage of 93.11% when combined.

YRGVYPFIY peptide giving the highest global coverage with a percentage of 54.79% of world population, a very low binding energy to HLA-A*68:01 of −25.1326 (Kcal/mol) and a strong interaction with HLA-C*07:02 with estimated IC50 of 35.88 making it one of the strongest T-cell candidates. Another strong peptide candidate is AVAAVSAAY peptide which had the best binding affinity of all five peptides to HLA-A*68:01 which is the most frequent MHC-I allele of −26.2570 (Kcal/mol) and a strong interaction with HLA-B*35:01 with an estimated IC50 of 12.76. HLYRGVYPF, RAAKFSHLY peptides which bind most to MHC class I alleles (10 alleles each).

As for MHC class II five peptides were selected as most potential peptides to actvate T-helper cell (CD4+) according to their population coverage globally, binding energy with the most frequent MHC-II allele HLA DRB1*01:01 and strength of interaction by the IC50 scores, HMIFASFIR, VFVVQKQLI, IAYPQLFNE, IFASFIRTA, and LRWAVSEAV (Global coverage of 95.03%, 91.03%, 93.63%, 92.82%, 79.58%, respectively). With an ideal total coverage of 99.70% of world population.

HMIFASFIR being one of the highly recommended peptides for giving the highest population coverage, binding affinity to HLA DRB1*01:01 of −9.356 (Kcal/mol) and IC50 (which indicated the strength of the interaction between the peptide and selected MHC allele) of 8.2 with HLA DRB5*01:01. LRWAVSEAV peptide had highest binding number to MHC II alleles (six alleles), a relatively high global population coverage (79.58%). Besides that, the peptide had a low binding energy required to bind to HLA-DRB1*01:01 (−18.8068 (Kcal/mol)) and strong interaction with HLA-DRB1*01:01 with and IC50 of 21. Another strong candidate MHC-II peptide is IFASFIRTA which shares few peptides with HMIFASFIR. The peptide had a high global population coverage estimated with 90.79%, binds to six MHC-II alleles, had a strong interaction with HLA-DQA1*01:02/DQB1*06:02 with an IC50 of 30.5, and gave the best docking result to HLA DRB1*01:01 with a binding energy of (−20.997 Kcal/mol)

Peptides binding to MHC class I and II alleles Combined were 61 peptides. The most important five peptides had the highest global population coverage percentage, HMIFASFIR, NFSHGSYEY, TETCAVAAV, VYKGRPLAI, and LRWAVSEAV (97.02%, 79.43%, 80.74%, 79.89%, and 80.56%, respectively). The five peptides altogether provided a total coverage of 99.85%.

Pyruvate kinase PK was selected for this study because it has proven to be a highly immunogenic protein in several organisms. A study by de Klerk N *et* al., 2012 has isolated PK and Frucose-bisphosphate Aldolase FBA as immunogens using flowcytometry from Madurella mycetomatis which causes Eumycetoma and it was expressed on its hyphae.^59^ A more recent study regarding M. Mycetomatis by Manofali A et al., 2018 confirmed used PK as an immunogenic for peptide-based vaccine design giving a moderately high global population coverage percentages in conclusion. Another study suggested, using N-terminal sequencing technique that Mycoplasma Synoviae and Mycoplasma Gallisepticum (major poultry pathogens) have 14 immunogenic proteins, among which, pyruvate kinase PK enzyme was by Bercic R. et al., 2007.^60^

In addition, various studies suggested PK to be an immunogenic stimulant and a useful target for vaccine design of candida albicans particularly. However, none of which actually conducted a peptide based study for PK in C. albicans. A study by Pitarch A. et al., 1999, using Gel electrohpresis, followed by western blotting technique for patients with systemic candidiasis detected over 18 immunogenic proteins in C. albicans, one of these was glycolytic enzymes including PK.^61^Another study by Mercedes Pardo et al, 2000. Using nanoelectrospray tandem mass spectrometry tech, suggested that C. albicans PK is an immunogenic protein.^62^ Other paper by swoboda et al. 1993, have suggested that pyruvate kinase protein gives strong interactions with IgM and IgG antibodies for candida albicans using pooled sera and concluded it being a proper antigen for candida albicans.^63^

This study was limited by being strictly computational, additionally, during analysis Calculations of some HLA class II alleles /epitopes (HLA DRB3*01:01, HLA DRB 5*01:01, HLA DPA 1*01, and HLA DRB 4*01:01) for population coverage were missed, that may influence in the accuracy of the population coverage results.

Many recent studies has proved the efficiency of peptide-based vaccines, moreover, few of them even made it to the clinical trials. A study by Matsumoto et al. 2016, reported a phase I study of a personalized peptide vaccine (PPV) for advanced urothelial carcinoma patients who failed treatment with MVAC. Ten patients with MVAC refractory advanced or metastatic urothelial cancer were treated twelve times using positive peptides chosen from 14 and 16 peptides in patients with HLA A24 and A2, respectively. The peptide vaccination was well tolerated with no major adverse effects and gave a notable rise in the IgG titer in patients.^64^ Another study by Schwartzentruber et al. 2011, had shown that testing two groups of patients with advanced melanoma. One group treated with interluken-2 only, and the second group treated with vaccine-interluken-2. The study concluded that patients of the second group response to the treatment was significantly improved than those of Group one.^65^

Peptide-based vaccination is a key role of combining a specific, desirable immune response, easy production, less laboring, with minimal immunological hazards, combining both T-cell and B-cell epitopes and could be used for both prevention or as a therapeutic tool.^66, 67^ peptide vaccine is poorly immunogenic when used alone, thus, a substance that facilitates and enhance antigen-antibody binding (adjuvant) is needed for better immune response.^68^ Future studies are required and highly recommended to subject these suggested peptides for clinical trials and select a suitable delivery system or adjuvant and strategies which could improve in-vivo and in-vitro T-cell and B-cell immune responses.

### Conclusion

Vaccine development for *C. albicans* is becoming a necessity due to, the growing resistance of the fungi to anti-fungal drugs, poor response to treatments and multiple recurrences to hospitalized patients. Peptide vaccines are well known for their efficiency, lack of hazard and less time consumption and that was the scope of this study. Fifteen peptides gave good population coverage percentages, in addition to strong interactions to MHC alleles of both classes I and II with varying degrees and poses of best possible binding energies to the most frequent peptides in the study HLA A*68:01 for MHC-I and HLA DRB1*01:01 or MHC-II. However, YRGVYPFIY, AVAAVSAAY, HMIFASFIR, LRWAVSEAV and IFASFIRTA T-cell peptides gave the best responses to the tests and therefore, are the most recommended peptides of this study for future studies in vitro and in vivo.

