## Supplementary material for "Multi-epitope Peptide Vaccine Prediction for Candida albicans targeting Pyruvate Kinase Protein; an Immunoinformatics Approach": table 3, figure 3, table 4, table 5, figure 4, figure 5

| No. | Residues | No. of residues | Score |
| --- | --- | --- | --- |
|  | B:S4, B:S5, B:L6, B:S7, B:W8, B:L9, B:S10, B:N11, B:F12, B:N13, B:V14, B:E15, B:T16, B:V17, B:P18, B:S19, B:K20, B:K361, B:A362, B:I363, B:A364, B:P366, B:Q367, B:F369, B:N370, B:E371, B:R373, B:S374, B:L375, B:A376, B:K377, B:K378, B:P379, B:T380, B:A381, B:T382, B:T385, B:C386, B:V388, B:A389, B:S392, B:A393, B:A394, B:Y395, B:E396, B:Q397, B:D398, B:A399, B:K400, B:Y418, B:V422, B:R429, B:Y448, B:D449, B:K450, B:P451, B:S452, B:I453, B:E454, B:N455, B:Q457, B:E458, B:D459, B:V460, B:E461, B:N462, B:R463, B:L464, B:R465, B:W466, B:V468, B:S469, B:E470, B:V472, B:E473, B:L474, B:G475, B:I476, B:I477, B:S478, B:K479, B:G480, B:D481, B:S482, B:I483, B:T485, B:Q487, B:G488, B:W489, B:T490, B:R491, B:G492, B:S493, B:G494, B:H495, B:S496, B:N497, B:T498, B:V499, B:R500, B:I501 | 101 | 0.758 |
| 2 | B:T96, B:G97, B:T98, B:T99, B:I100, B:G101, B:D102, B:K103, B:D104, B:Y105, B:P106, B:I107, B:P108, B:P109, B:N110, B:H111, B:E112, B:M113, B:I114, B:F115, B:T116, B:T117, B:D118, B:D119, B:A120, B:Y121, B:K122, B:T123, B:K124, B:C125, B:D126, B:D127, B:K128, B:V129, B:M130, B:Y131, B:I132, B:Y134, B:K135, B:N136, B:I137, B:T138, B:K139, B:V140, B:I141, B:A142, B:P143, B:G144, B:K145, B:I146, B:V153, B:L154, B:S155, B:F156, B:E157, B:V158, B:I159, B:S160, B:V161, B:D162, B:D163, B:E164, B:Q165, B:T166, B:L167, B:K168, B:V169, B:R170, B:S171, B:L172, B:N173, B:A174, B:G175, B:K176, B:I177, B:S178, B:S179, B:H180, B:K181, B:L185, B:P186, B:G187, B:T188, B:D189, B:V190, B:D191, B:L192 | 87 | 0.755 |
| 3 | B:G233, B:E234, B:E235 | 3 | 0.722 |
| 4 | B:G32, B:P33, B:K34, B:T35, B:N36, B:N37, B:V38, B:D39, B:V40, B:V42, B:K43, B:K46, B:Q65, B:S66, B:D69, B:N70, B:R72, B:K73, B:S74, B:E76, B:V77, B:Y78, B:A340 | 23 | 0.648 |
| 5 | B:S307, B:N311, B:P312, B:R313, B:P314, B:T315, B:R316, B:A317, B:G337, B:K341, B:G342, B:N343, B:Y344, B:V346, B:E347, B:M351 | 16 | 0.63 |
| 6 | B:K208, B:N209, B:K210 | 3 | 0.598 |
| 7 | B:G59, B:S60, B:Y61, B:E62, B:Y63 | 5 | 0.577 |
| 8 | B:I200, B:A201, B:K204, B:V231 | 4 | 0.572 |
| 9 | B:T221, B:N223, B:L226 | 3 | 0.535 |

**Table3**. Full List of Predicted discontinuous B-cell epitopes with number of Residues and their scores using ElliPro prediction tool

| 10 | D:S4, D:S5, D:L6, D:S7, D:W8, D:L9, D:S10, D:N11, D:F12, D:N13, D:V14, D:E15, D:T16, D:V17, D:P18, D:S19, D:K20, D:K361, D:A362, D:I363, D:A364, D:P366, D:Q367, D:F369, D:N370, D:E371, D:R373, D:S374, D:L375, D:A376, D:K377, D:K378, D:P379, D:T380, D:A381, D:T382, D:T385, D:C386, D:V388, D:A389, D:S392, D:A393, D:A394, D:Y395, D:E396, D:Q397, D:D398, D:A399, D:K400, D:Y418, D:V422, D:R429, D:Y448, D:D449, D:K450, D:P451, D:S452, D:I453, D:E454, D:N455, D:Q457, D:E458, D:D459, D:V460, D:E461, D:N462, D:R463, D:L464, D:R465, D:W466, D:V468, D:S469, D:E470, D:V472, D:E473, D:L474, D:G475, D:I476, D:I477, D:S478, D:K479, D:G480, D:D481, D:S482, D:I483, D:T485, D:Q487, D:G488, D:W489, D:T490, D:R491, D:G492, D:S493, D:G494, D:H495, D:S496, D:N497, D:T498, D:V499, D:R500, D:I501 | 101 | 0.758 |
| --- | --- | --- | --- |
| 11 | D:T96, D:G97, D:T98, D:T99, D:I100, D:G101, D:D102, D:K103, D:D104, D:Y105, D:P106, D:I107, D:P108, D:P109, D:N110, D:H111, D:E112, D:M113, D:I114, D:F115, D:T116, D:T117, D:D118, D:D119, D:A120, D:Y121, D:K122, D:T123, D:K124, D:C125, D:D126, D:D127, D:K128, D:V129, D:M130, D:Y131, D:I132, D:Y134, D:K135, D:N136, D:I137, D:T138, D:K139, D:V140, D:I141, D:A142, D:P143, D:G144, D:K145, D:I146, D:V153, D:L154, D:S155, D:F156, D:E157, D:V158, D:I159, D:S160, D:V161, D:D162, D:D163, D:E164, D:Q165, D:T166, D:L167, D:K168, D:V169, D:R170, D:S171, D:L172, D:N173, D:A174, D:G175, D:K176, D:I177, D:S178, D:S179, D:H180, D:K181, D:L185, D:P186, D:G187, D:T188, D:D189, D:V190, D:D191, D:L192 | 87 | 0.755 |
| 12 | D:G233, D:E234, D:E235 | 3 | 0.722 |
| 13 | D:G32, D:P33, D:K34, D:T35, D:N36, D:N37, D:V38, D:D39, D:V40, D:V42, D:K43, D:K46, D:Q65, D:S66, D:D69, D:N70, D:R72, D:K73, D:S74, D:E76, D:V77, D:Y78, D:A340 | 23 | 0.648 |
| 14 | D:S307, D:N311, D:P312, D:R313, D:P314, D:T315, D:R316, D:A317, D:G337, D:K341, D:G342, D:N343, D:Y344, D:V346, D:E347, D:M351 | 16 | 0.63 |
| 15 | D:K208, D:N209, D:K210 | 3 | 0.598 |
| 16 | D:G59, D:S60, D:Y61, D:E62, D:Y63 | 5 | 0.577 |
| 17 | D:I200, D:A201, D:K204, D:V231 | 4 | 0.572 |
| 18 | D:T221, D:N223, D:L226 | 3 | 0.535 |

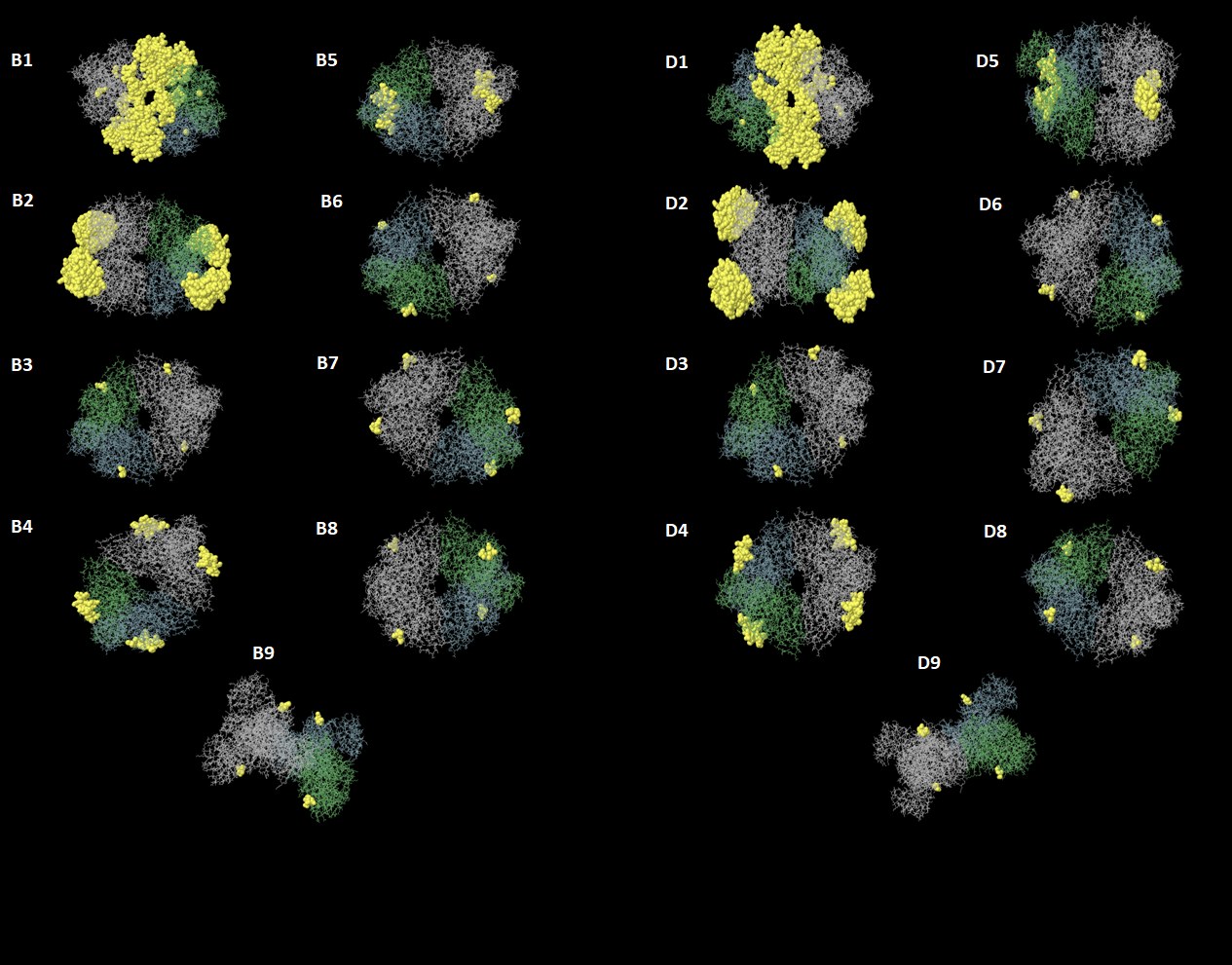

**Figure 3:** Three-dimensional representation of non-linear epitopes (B1–B9, D1-D9) of the highly immunogenic pyruvate Kinase PK protein of *C. albicans* using ElliPro prediction tool. The epitopes are highlighted in yellow surface, and the bulk of PK protein is depicted in grey.

**Table 4**: List of most promising epitopes that had a good binding affinity with MHC-I alleles in terms of IC50 and Percentile rank.

| Peptide | Start | End | Allele | ic50 | Percentile Rank |
| --- | --- | --- | --- | --- | --- |
| AVAAVSAAY* | 387 | 395 | HLA-A*03:01 | 363.95 | 1.5 |
|  | 387 | 395 | HLA-A*11:01 | 102.38 | 0.69 |
|  | 387 | 395 | HLA-A*26:01 | 45.86 | 0.05 |
|  | 387 | 395 | HLA-A*29:02 | 26.57 | 0.2 |
|  | 387 | 395 | HLA-A*30:02 | 15 | 0.03 |
|  | 387 | 395 | HLA-B*15:01 | 19.77 | 0.11 |
|  | 387 | 395 | HLA-B*15:02 | 155.4 | 0.07 |
|  | 387 | 395 | HLA-B*35:01 | 12.76 | 0.06 |
| HLYRGVYPF* | 438 | 446 | HLA-A*02:06 | 272.9 | 1.8 |
|  | 438 | 446 | HLA-A*23:01 | 88.52 | 0.29 |
|  | 438 | 446 | HLA-A*29:02 | 217.62 | 0.85 |
|  | 438 | 446 | HLA-A*32:01 | 7.03 | 0.02 |
|  | 438 | 446 | HLA-B*15:01 | 20.19 | 0.11 |
|  | 438 | 446 | HLA-B*15:02 | 22.64 | 0.02 |
|  | 438 | 446 | HLA-B*35:01 | 82.9 | 0.27 |
|  | 438 | 446 | HLA-C*07:02 | 99.78 | 0.03 |
|  | 438 | 446 | HLA-C*12:03 | 69.01 | 0.15 |
|  | 438 | 446 | HLA-C*14:02 | 145.33 | 0.23 |
| RAAKFSHLY* | 432 | 440 | HLA-A*11:01 | 259.26 | 1.6 |
|  | 432 | 440 | HLA-A*29:02 | 65.75 | 0.4 |
|  | 432 | 440 | HLA-A*30:02 | 14.79 | 0.02 |
|  | 432 | 440 | HLA-A*68:01 | 485.71 | 2.2 |
|  | 432 | 440 | HLA-B*15:01 | 183.28 | 0.82 |
|  | 432 | 440 | HLA-B*35:01 | 129.49 | 0.34 |
|  | 432 | 440 | HLA-B*57:01 | 40.73 | 0.16 |
|  | 432 | 440 | HLA-B*58:01 | 16.15 | 0.09 |
|  | 432 | 440 | HLA-C*12:03 | 48.21 | 0.11 |
|  | 432 | 440 | HLA-C*15:02 | 399.89 | 0.21 |
| YRGVYPFIY* | 440 | 448 | HLA-A*29:02 | 90.75 | 0.51 |
|  | 440 | 448 | HLA-B*27:05 | 299.39 | 1.2 |
|  | 440 | 448 | HLA-C*06:02 | 332.17 | 0.12 |
|  | 440 | 448 | HLA-C*07:01 | 484.79 | 0.14 |
|  | 440 | 448 | HLA-C*07:02 | 35.88 | 0.02 |
| YVDDGVLSF* | 148 | 156 | HLA-A*01:01 | 360.12 | 0.52 |
|  | 148 | 156 | HLA-A*02:06 | 27.1 | 0.32 |
|  | 148 | 156 | HLA-A*29:02 | 161.46 | 0.72 |
|  | 148 | 156 | HLA-B*15:01 | 427.25 | 1.6 |
|  | 148 | 156 | HLA-B*35:01 | 23.88 | 0.1 |
|  | 148 | 156 | HLA-C*03:03 | 113.92 | 0.34 |
|  | 148 | 156 | HLA-C*05:01 | 3.9 | 0.01 |
|  | 148 | 156 | HLA-C*12:03 | 10.26 | 0.03 |
|  | 148 | 156 | HLA-C*14:02 | 112.6 | 0.2 |

**Table 5:** List of the five promising core sequence epitopes that had a strong binding affinity with MHC-II in terms of IC50 and Percentile Ranks

| Core peptide Sequence | Start | End | Allele | IC50 | Rank |
| --- | --- | --- | --- | --- | --- |
| HMIFASFIR | 208 | 222 | HLA-DPA1*01/DPB1*04:01 | 72.3 | 4.14 |
|  | 209 | 223 | HLA-DPA1*01/DPB1*04:01 | 74.2 | 4.22 |
|  | 210 | 224 | HLA-DPA1*01/DPB1*04:01 | 75.1 | 4.26 |
|  | 208 | 222 | HLA-DPA1*01:03/DPB1*02:01 | 69 | 6.49 |
|  | 210 | 224 | HLA-DPA1*01:03/DPB1*02:01 | 72.6 | 6.72 |
|  | 207 | 221 | HLA-DPA1*02:01/DPB1*01:01 | 92.9 | 9.77 |
|  | 207 | 221 | HLA-DPA1*02:01/DPB1*05:01 | 57.3 | 0.93 |
|  | 208 | 222 | HLA-DPA1*02:01/DPB1*05:01 | 64.6 | 1.11 |
|  | 206 | 220 | HLA-DPA1*02:01/DPB1*05:01 | 68.6 | 1.21 |
|  | 209 | 223 | HLA-DRB5*01:01 | 8.2 | 1.83 |
|  | 212 | 226 | HLA-DRB5*01:01 | 63 | 11.62 |
| IAYPQLFNE | 359 | 373 | HLA-DPA1*01:03/DPB1*02:01 | 56.9 | 5.65 |
|  | 360 | 374 | HLA-DPA1*01:03/DPB1*02:01 | 59.1 | 5.81 |
|  | 358 | 372 | HLA-DPA1*01:03/DPB1*02:01 | 60.8 | 5.93 |
|  | 361 | 375 | HLA-DPA1*01:03/DPB1*02:01 | 73.6 | 6.79 |
|  | 360 | 374 | HLA-DPA1*02:01/DPB1*01:01 | 28.2 | 2.63 |
|  | 359 | 373 | HLA-DPA1*02:01/DPB1*01:01 | 29.6 | 2.81 |
|  | 361 | 375 | HLA-DPA1*02:01/DPB1*01:01 | 38.8 | 3.95 |
|  | 358 | 372 | HLA-DPA1*02:01/DPB1*01:01 | 39.3 | 4.01 |
|  | 360 | 374 | HLA-DPA1*03:01/DPB1*04:02 | 43.5 | 5.1 |
| IFASFIRTA | 209 | 223 | HLA-DPA1*01:03/DPB1*02:01 | 67.7 | 6.4 |
|  | 211 | 225 | HLA-DPA1*01:03/DPB1*02:01 | 87.8 | 7.66 |
|  | 209 | 223 | HLA-DPA1*02:01/DPB1*01:01 | 41.5 | 4.28 |
|  | 210 | 224 | HLA-DPA1*02:01/DPB1*01:01 | 45.6 | 4.76 |
|  | 208 | 222 | HLA-DPA1*02:01/DPB1*01:01 | 46.6 | 4.88 |
|  | 211 | 225 | HLA-DPA1*02:01/DPB1*01:01 | 54.1 | 5.76 |
|  | 212 | 226 | HLA-DPA1*02:01/DPB1*01:01 | 91.8 | 9.66 |
|  | 211 | 225 | HLA-DQA1*01:02/DQB1*06:02 | 30.5 | 1.31 |
|  | 210 | 224 | HLA-DQA1*01:02/DQB1*06:02 | 32.5 | 1.46 |
|  | 212 | 226 | HLA-DQA1*01:02/DQB1*06:02 | 39.1 | 2.01 |
|  | 213 | 227 | HLA-DQA1*01:02/DQB1*06:02 | 48.4 | 2.81 |
|  | 214 | 228 | HLA-DQA1*01:02/DQB1*06:02 | 74.8 | 5.06 |
| VFVVQKQLI | 277 | 291 | HLA-DPA1*01:03/DPB1*02:01 | 91.7 | 7.89 |
|  | 276 | 290 | HLA-DPA1*02:01/DPB1*01:01 | 66.1 | 7.08 |
|  | 277 | 291 | HLA-DPA1*02:01/DPB1*01:01 | 66.5 | 7.12 |
|  | 278 | 292 | HLA-DPA1*02:01/DPB1*01:01 | 72.2 | 7.74 |
|  | 275 | 289 | HLA-DPA1*02:01/DPB1*01:01 | 79.8 | 8.52 |
|  | 279 | 293 | HLA-DPA1*02:01/DPB1*01:01 | 84.8 | 9 |
|  | 275 | 289 | HLA-DRB1*07:01 | 73.7 | 11.55 |
|  | 274 | 288 | HLA-DRB1*07:01 | 81.9 | 12.4 |
|  | 278 | 292 | HLA-DRB4*01:01 | 15.5 | 0.67 |
|  | 277 | 291 | HLA-DRB4*01:01 | 18.4 | 0.9 |
|  | 279 | 293 | HLA-DRB4*01:01 | 20.5 | 1.08 |
|  | 276 | 290 | HLA-DRB4*01:01 | 23.9 | 1.38 |
|  | 280 | 294 | HLA-DRB4*01:01 | 31.4 | 2.04 |
|  | 275 | 289 | HLA-DRB4*01:01 | 35.1 | 2.37 |
|  | 274 | 288 | HLA-DRB4*01:01 | 45.1 | 3.27 |
| LRWAVSEAV | 464 | 478 | HLA-DQA1*05:01/DQB1*03:01 | 35.3 | 6.64 |
|  | 463 | 477 | HLA-DQA1*05:01/DQB1*03:01 | 37.2 | 6.97 |
|  | 460 | 474 | HLA-DQA1*05:01/DQB1*03:01 | 38.3 | 7.15 |
|  | 461 | 475 | HLA-DQA1*05:01/DQB1*03:01 | 39.6 | 7.37 |
|  | 462 | 476 | HLA-DQA1*05:01/DQB1*03:01 | 44.3 | 8.12 |
|  | 459 | 473 | HLA-DQA1*05:01/DQB1*03:01 | 48.4 | 8.74 |
|  | 458 | 472 | HLA-DQA1*05:01/DQB1*03:01 | 60.8 | 10.45 |
|  | 461 | 475 | HLA-DRB1*01:01 | 21 | 11.79 |
|  | 460 | 474 | HLA-DRB1*01:01 | 30.8 | 15.75 |
|  | 461 | 475 | HLA-DRB1*03:01 | 98.4 | 4.96 |
|  | 460 | 474 | HLA-DRB1*03:01 | 99.3 | 4.98 |
|  | 458 | 472 | HLA-DRB1*07:01 | 24.1 | 4.65 |
|  | 459 | 473 | HLA-DRB1*07:01 | 27.1 | 5.21 |
|  | 459 | 473 | HLA-DRB1*09:01 | 39.3 | 2.35 |
|  | 458 | 472 | HLA-DRB1*09:01 | 70.1 | 4.77 |
|  | 460 | 474 | HLA-DRB1*13:02 | 64 | 4.29 |
|  | 461 | 475 | HLA-DRB1*13:02 | 65.1 | 4.35 |
|  | 462 | 476 | HLA-DRB1*13:02 | 95.7 | 5.87 |

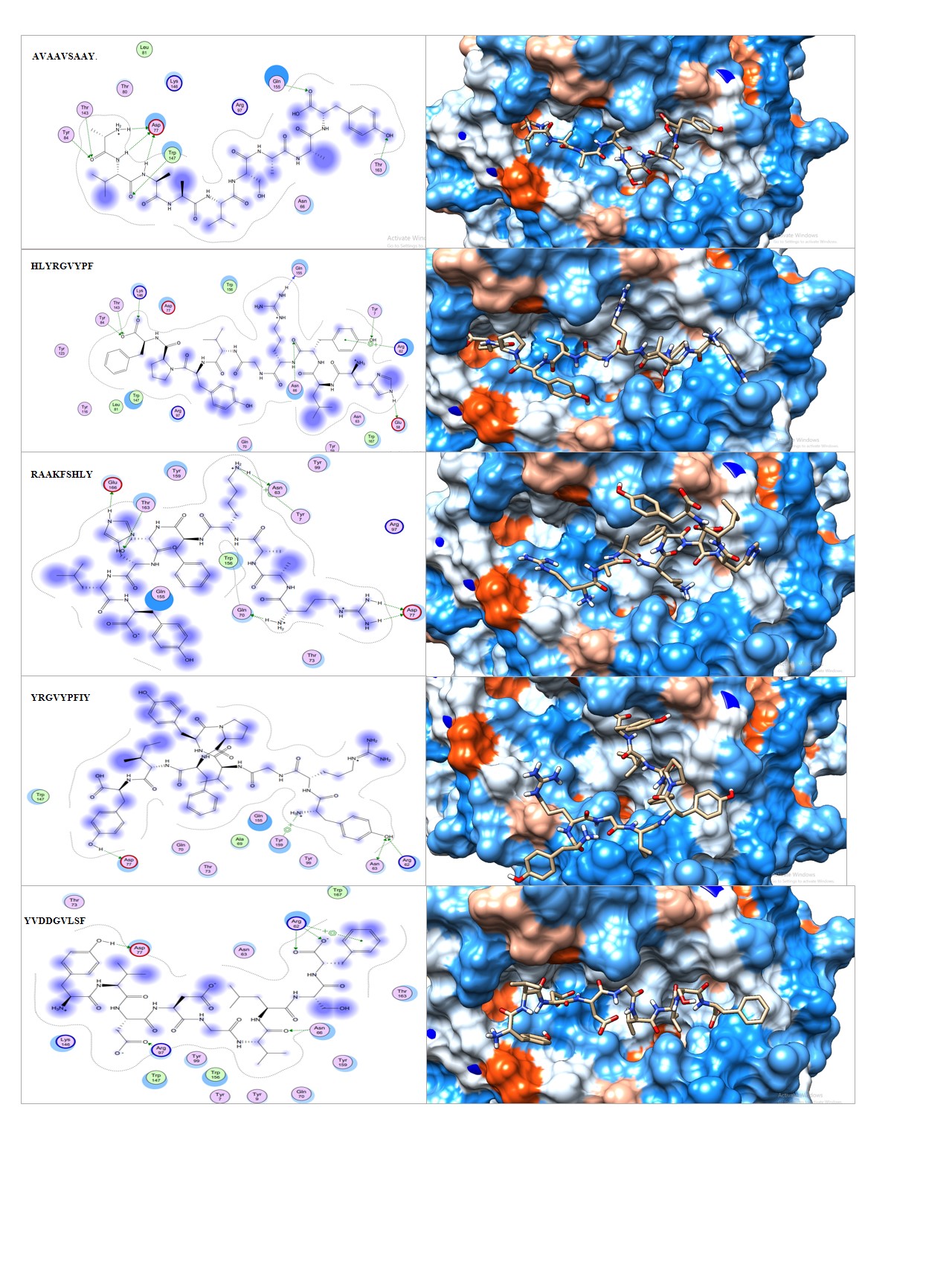

**Figure 4.** Illustrates 2 and 3-dimentional interaction of the best docking poses with HLA-A*68:01 MHC-I allele for five promising peptides **A.** AVAAVSAAY **B.** HLYRGVYPF **C.** RAAKFSHLY **D.** YRGVYPFIY **and E.** YVDDGVLSF.

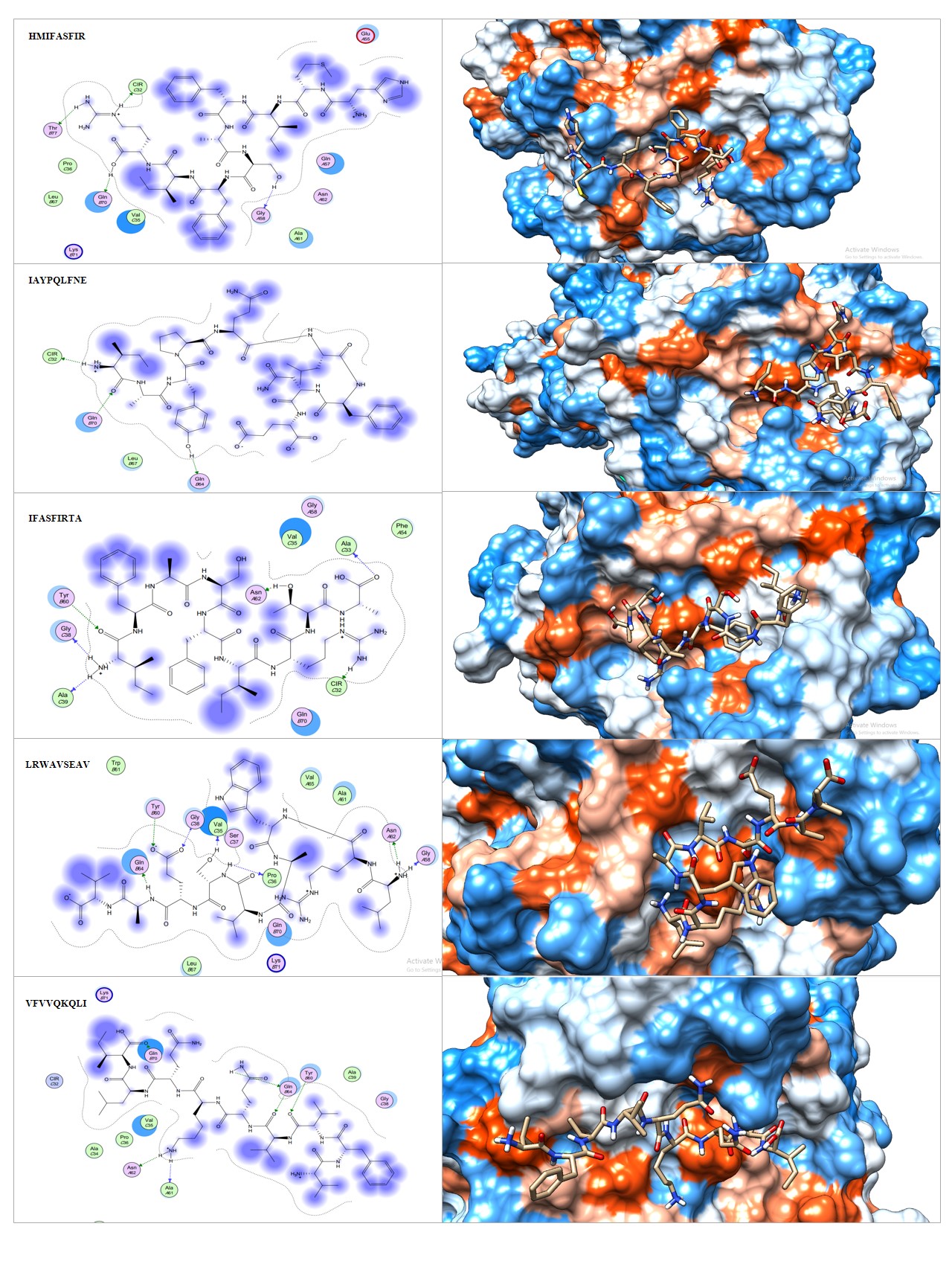
 **Figure 5.** Illustrates 2 and 3-dimentional interaction of the best docking poses with HLA-DRB1*01:01 MHC-II allele for five promising peptides **A.** HMIFASFIR **B.** IAYPQLFNE **C.** IFASFIRTA **D.** LRWAVSEAV **and E.** VFVVQKQLI
